# Systematic analysis of protein targets associated with adverse events of drugs from clinical trials and post-marketing reports

**DOI:** 10.1101/2020.06.12.135939

**Authors:** Ines A. Smit, Avid M. Afzal, Chad H. G. Allen, Fredrik Svensson, Thierry Hanser, Andreas Bender

## Abstract

Adverse drug reactions (ADRs) are undesired effects of medicines that can harm patients and are a significant source of attrition in drug development. ADRs are anticipated by routinely screening drugs against secondary pharmacology protein panels. However, there is still a lack of quantitative information on the links between these off-target proteins and the risk of ADRs in humans. Here, we present a systematic analysis of associations between measured and predicted *in vitro* bioactivities of drugs, and adverse events (AEs) in humans from two sources of data: the Side Effect Resource (SIDER), derived from clinical trials, and the Food and Drug Administration Adverse Event Reporting System (FAERS), derived from post-marketing surveillance. The ratio of a drug’s *in vitro* potency against a given protein relative to its therapeutic unbound drug plasma concentration was used to select proteins most likely to be relevant to *in vivo* effects. In examining individual target bioactivities as predictors of AEs, we found a trade-off between the Positive Predictive Value and the fraction of drugs with AEs that can be detected, however considering sets of multiple targets for the same AE can help identify a greater fraction of AE-associated drugs. Of the 45 targets with statistically significant associations to AEs, 30 are included on existing safety target panels. The remaining 15 targets include 8 carbonic anhydrases, of which CA5B was significantly associated with cholestatic jaundice. We include the full quantitative data on associations between *in vitro* bioactivities and AEs in humans in this work, which can be used to make a more informed selection of safety profiling targets.

## Introduction

Adverse drug reactions (ADRs) are a major cause of harm to patients, including hospitalization, disability, and mortality *(1)*. Safety issues in preclinical and clinical studies are a major obstacle in the development of new drugs *(2)*, whereas post-marketing withdrawals have also occurred, e.g. fenfluramine was withdrawn in 1997 due to associated cardiac valvulopathy related to agonism at the serotonin 2B receptor (5-HT_2B_) *(3, 4)*.

Up to 75% of ADRs occurring in hospitals or leading to hospital admissions have been estimated to be dose-dependent and predictable from the drug’s pharmacology, i.e. defined drug-protein interactions *(5)*. Thus, to anticipate ADRs in early drug discovery, pharmaceutical companies commonly screen drug candidates for binding against a panel of safety targets. One such panel published by Bowes et al. consists of 44 proteins which were compiled as the consensus set of targets screened by major pharmaceutical companies and it includes G protein-coupled receptors (GPCR), ion channels, enzymes, transporters and nuclear receptors *(2)*. Whitebread et al. compiled a set of 36 targets related to cardiovascular toxicity *(6)*, while a panel published by Lynch et al. includes over 70 targets *(7)*.

The above-mentioned safety target panels share between ~30-80% of targets, showing that there are differences in targets chosen for screening. Authors from the U.S. Food and Drug Administration (FDA) stated that “the panels of targets that are employed [for regulatory submissions] vary widely and are often selected without justification or a description of their relevance to human safety” *(8)*. This is related to the general lack of *quantitative* information on the risk associated with safety targets, since systematic information about their predictivity is rarely available *(7, 9)*. This limits the conclusions one can draw currently from secondary pharmacology screens, even when a comprehensive set of targets is screened, because in many cases we cannot sufficiently extrapolate to *in vivo* effects in humans.

Some previous studies have tried to address the lack of evidence on safety targets and their relationships to ADRs observed in humans. Lynch et al. performed an extensive literature survey to back up the evidence behind the targets in the Abbvie panel, although the relationships were not quantified *(7)*. Another, more quantitative approach was used by Kuhn et al. in their statistical analysis of side effects listed on drug package inserts from the Side Effect Resource (SIDER) and *in vitro* bioactivity data from the STITCH3 database *(10)*. For 732 out of the 1,428 (51%) side effects analysed, the authors were able to identify statistically overrepresented targets *(10)*. These proteins provided a plausible causal explanation for the side effect in over 70% of the drug-side effect pairs based on supporting literature evidence *(10)*. A similar study by Duran-Frigola and Aloy found that 41% of studied side effects from SIDER could be statistically related to 79 therapeutic drug targets from Drugbank *(11)*. Both studies have the limitation that quantifications of the associations beyond the p-value were not provided, which is important for estimating the effect size of associations. Krejsa et al. did provide Positive Predictive Values in their study, e.g. 50% of drugs with an IC50 between 0.1-1 μM at the muscarinic receptor M1 caused tachycardia, based on the Bioprint database *(9)*. However, data for only three such target-adverse event (AE) combinations were reported *(9)*. In a recent study, Deaton et al. used pharmacological and genetic data to select targets for screening, but their quantification focused on the correspondence between pharmacological effects and genetic phenotypes related to targets *(12)*.

To the best knowledge of the authors, no current study systematically addresses links between protein target activities and observed ADRs while taking into account drug pharmacokinetics on a large scale. While this is indeed more difficult due to scarcity of available data, we can only expect to observe effects of drugs *in vivo* when sufficiently high drug concentrations at the target site are reached *(13, 14)*. In a case study of risperidone, Maciejewski et al. showed that the integration of the unbound plasma concentration C_max_ was crucial for rationalizing the relationship between the dopamine D2 receptor and galactorrhoea *(15)*. While concentrations at the target site depend on complex factors such as drug tissue distributions and subcellular target locations, blood plasma concentrations are commonly used as an approximation in practice *(2, 13)*, enabling the *in vitro* potency of drugs to be related to their relevance to *in vivo* effects.

We hence compiled a set of unbound plasma concentrations from a number of publications and the ChEMBL database, and used these to identify target-AE associations between *in vitro* bioactivity and AEs in humans, according to the workflow in Fig. 1.

**Fig. 1.**
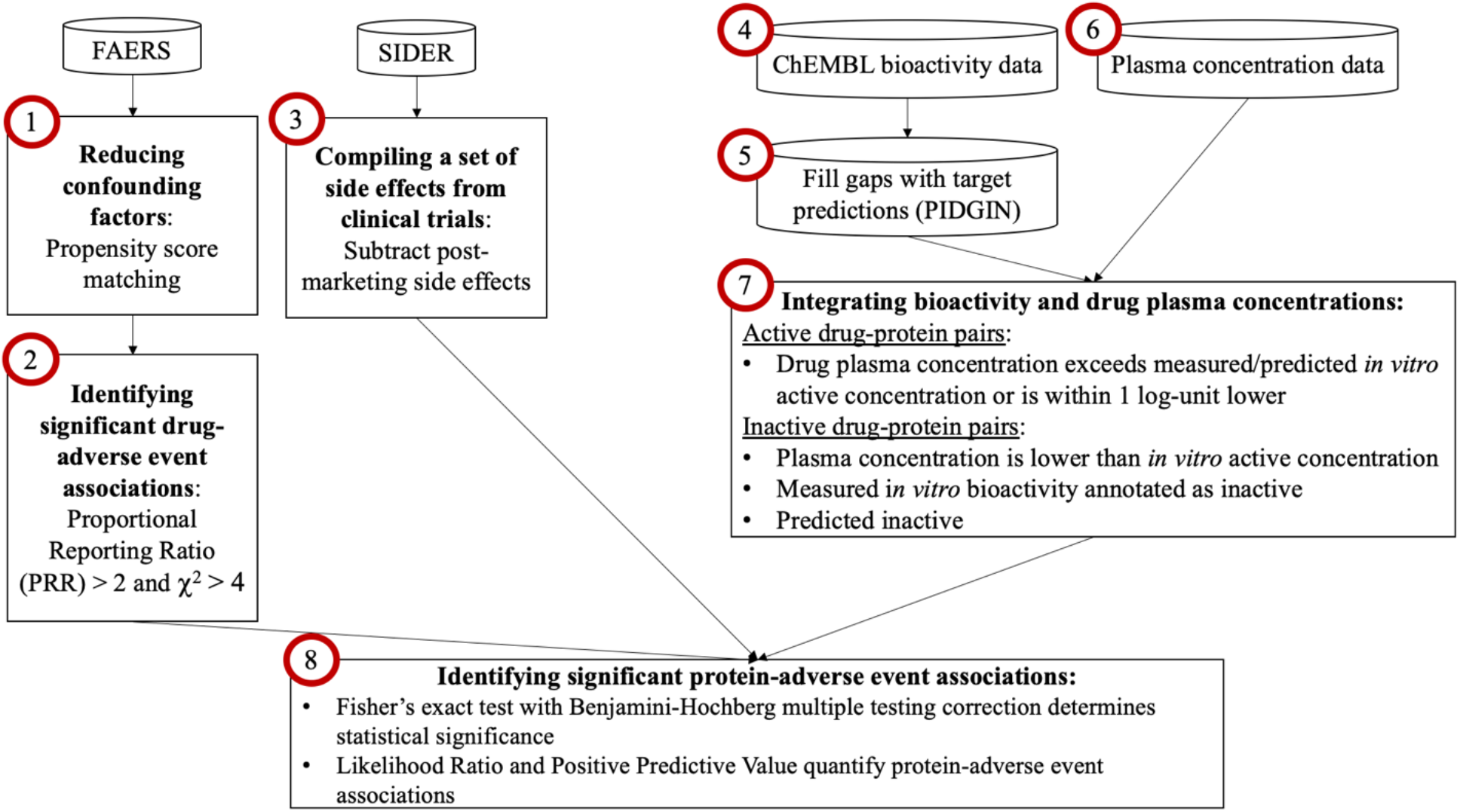
Overview of the workflow of the study. Each numbered item is described in more details in the Materials & Methods.

In this work, we analysed data from the FDA Adverse Event Reporting System (FAERS), which contains post-marketing reports of AEs submitted by health care practitioners, consumers and drug manufacturers *(16)*. We used the curated AEOLUS version of FAERS, covering the years 2004-2015 and containing close to five million unique case reports *(17)*. Since FAERS data is derived from large samples of diverse patients and potentially long-term use of medicines, it may contain ADRs that would be missed during clinical trials *(18)*. However, various biases can affect FAERS, such as sampling and reporting biases *(18, 19)*. To counteract these, we applied a previously reported method, Propensity Score Matching (PSM), which successfully reduced the effects of confounding factors such as drug indications, concomitant medications, sex, and age in FAERS in a previous study *(19)*. Subsequently, we identified drug-AE associations using the Proportional Reporting Ratio (PRR) (Fig. 1, steps 1 and 2).

Post-marketing reporting systems are focused on serious and unexpected AEs, thus to compile more complete AE profiles we also used the SIDER dataset, which is based on side effects observed during clinical trials that were summarised and subsequently extracted from drug package inserts by text-mining *(20)*. Clinical trials often include a control group and thus may be less biased than post-marketing data, but their sample size and patient population is more limited *(20)*. To focus exclusively on the clinical effects, we subtracted the small amount of post-marketing effects present in SIDER (Fig. 1, step 3). Both FAERS and SIDER hence provided us with drug-AE associations for further analysis.

Next, we used the *in vitro* bioactivity data from the ChEMBL database and supplemented this with ligand-based target predictions, which was necessary due to the known sparsity of ChEMBL data (Fig. 1, steps 4 and 5) *(21, 22)*. This step hence provided us with drug-target associations. We finally compared the compiled unbound plasma concentrations with the measured or predicted *in vitro* bioactivities to identify those target interactions that could be responsible for AEs, given pharmacokinetic properties.

This analysis thus provides a large-scale analysis of links between drugs, their protein targets, and AEs in a quantitative manner, while taking into account plasma drug exposure.

## Results

### Overall dataset analysis

The set of statistically significant drug-AE relationships contains 1,263 unique drugs related to 3,365 unique AEs – counting Preferred Terms (PT), the main AE terms of the Medical Dictionary for Regulatory Activities (MedDRA) vocabulary – based on FAERS (Data File S 1). The respective set based on SIDER, restricted to events from clinical trials, contains 1,027 unique drugs and 1,130 unique AEs. A total of 696 drugs overlap between the datasets. In terms of the drugs’ distribution across Anatomical Therapeutic Chemical (ATC) classes, representing drug indications, both datasets are largely similar to the set of all small molecule drugs in ChEMBL (Fig. S 1). Despite some differences, such as the FAERS and SIDER datasets containing 17% more drugs used for nervous system indications than the set of all marketed small molecules, both datasets represent all ATC classes of marketed drugs (Fig. S 1). The distribution of AEs across System Organ Classes (SOC), the highest level of the MedDRA hierarchy, is shown in Fig. S 2. ‘Investigations’ are the largest group of AEs in FAERS, which is a highly diverse SOC that includes events such as electrocardiogram observations and changes in blood pressure, whereas nervous system disorders are the largest group in the SIDER dataset (Fig. S 2). Compared to SIDER, FAERS contains a higher diversity of events in ‘injury, poisoning and procedural complications’, neoplasms, and ‘surgical and medical procedures’ (Fig. S 2). These differences are consistent with the more diverse, post-marketing origin of FAERS that also includes reports of accidents *(16)*. SIDER contains more diverse events in ‘eye disorders’ and ‘skin and subcutaneous tissue disorders’, suggesting these events are more typically reported during clinical trials (Fig. S 2). The bioactivity data from ChEMBL has a matrix density of 5% of measured active and inactive datapoints, with the rest of the data being missing (Data File S 2). Using target prediction to supplement the measurements, while using the model performance and applicability domain thresholds described in the Methods, the matrix density increased to 25% (Data File S 2). We retrieved the unbound plasma concentrations for 466 drugs from the FAERS and SIDER datasets, which have a median concentration of pMolar 6.6 and a standard deviation (STD) of 1.5 (Fig. S 3). As expected, the unbound concentrations are lower than the total plasma concentrations, which have a median concentration of pMolar 5.8 (Fig. S 3). Using the ratio of the pXC50 and plasma concentration to assign the ‘activity call’ resulted in 3.6% of measured drug-target datapoints being assigned as active, while this is 0.1% for predicted datapoints (Data File S 2). Thus, the target predictions primarily added inactive datapoints.

After overlapping the three data types (AE, bioactivity, and plasma concentrations), we use the drug-AE and drug-target relationships to infer target-AE relationships that are potentially responsible for observed AEs. Based on FAERS data we analysed 197,236 target-AE combinations (100 unique targets and 3,278 unique AEs, Data File S 3), and based on SIDER a total of 42,652 target-AE pairs (79 unique targets and 982 unique AEs, Data File S 4). Hence, the FAERS dataset contains about four times as many target-AE pairs as the SIDER one.

### Impact of using plasma concentrations on target-AE associations

The above numbers are based on incorporating plasma concentrations to derive active drug-protein pairs. To investigate the effect of using the plasma concentrations, we also created a baseline dataset using a constant bioactivity cut-off of pChEMBL ≥ 6 (1 μM) for comparison. This dataset contains 313,661 target-AE pairs (182 unique targets and 3,340 unique AEs, Data File S 5) based on FAERS and 89,105 target-AE pairs (167 unique targets and 1,119 unique AEs, Data File S 6) based on SIDER. Thus, this dataset is roughly twice as large as the former dataset using the plasma concentrations in terms of the number of datapoints and unique targets.

Comparing the distribution of datapoints across target classes, shows that using plasma concentrations together with requiring a minimum of 5 datapoints per target and AE results in more GPCR datapoints being retained, the fraction of which increased from ~30% to ~48% of datapoints, and fewer kinase datapoints, which decreased from ~22% to ~5%, compared to using the constant cut-off (Fig. S 4). This can be explained by few pXC50 datapoints for the same kinase being available, resulting in kinases being disproportionately affected by the minimum of 5 datapoints per target when the available data is halved due to requiring plasma concentrations.

To examine the impact of using plasma concentrations on deriving target-AE associations, we compared the statistically significant (corrected p-value ≤ 0.05) associations found when using the constant pChEMBL cut-off to derive active drug-protein pairs versus taking unbound concentrations into account for this purpose. This shows that 41% target-AE associations found when using unbound concentrations are not significant when using the constant cut-off. Thus, the significant target-AE associations found when using plasma concentrations is not a direct subset of the dataset using a constant cut-off, showing that using plasma concentrations results in a different set of target-AE associations. With regards to the accuracy of the associations, the lack of a gold standard of target-AE associations makes it difficult to evaluate both outcomes *(8)*.

Therefore we chose to evaluate the two datasets by comparing them to three previously reported safety target panels *(2, 6, 7)*. The overall retrieval of these previously reported associations, based on overlapping MedDRA High Level Terms (HLTs) - the hierarchy level above the MedDRA PT - as statistically significant in our study is 6% using the unbound concentrations and 12% when using the absolute cut-off. However, while the retrieval is lower when using plasma concentrations, those target-AE pairs that are retrieved have larger Likelihood Ratios (LR), indicating greater strength of association (Fig. 2). Similarly, the Positive Predictive Value (PPV) and value-added PPV, i.e. the PPV minus the prevalence *(23)*, of the associations that are retrieved are higher when using unbound plasma concentrations (Fig. 2). This indicates that using plasma concentrations has fewer false positive signals compared to using the constant cut-off. However, the fraction of drugs associated with the AE per target, which would be the ‘detection’ rate of AE-associated drugs if the target is used as a predictor of AEs, is lower for the target-AE associations derived from using plasma concentrations (Fig. 2). This indicates that the greater LRs and precision when using plasma concentrations is at the cost of lower recall. One interpretation of these results is that using plasma concentrations retrieves known signals more strongly and precisely, but that recall could be limited by the amount of data available. Despite the lower overall recall, we noticed that some familiar examples, such as hERG-torsade de pointes and 5-HT_2B_-cardiotoxicity in the FAERS dataset, as well as dopamine D2-galactorrhoea in both the FAERS and SIDER datasets, were only significant when using plasma concentrations and not when using the constant cut-off. Since the plasma concentrations are a novel aspect of the current study and the results suggest they retrieve known signals with greater LRs and PPVs, for the remainder of this work we will focus on the results based on using plasma concentrations.

**Fig. 2.**
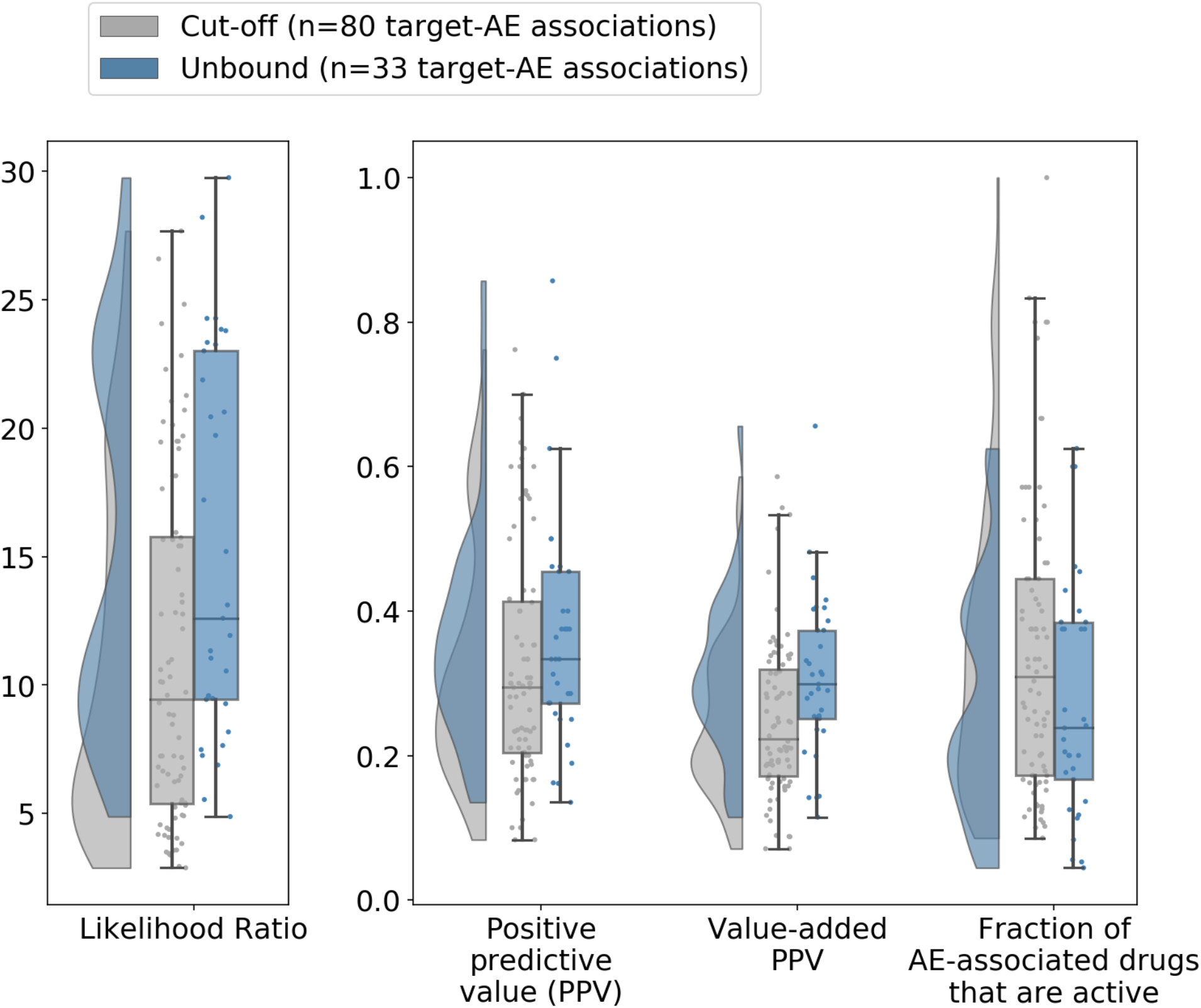
Quantification of target-AE associations reported in previous studies that were retrieved as significant in the current work when using a constant bioactivity threshold (pChEMBL ɥ 6) to define active drug-target pairs, versus using the ratio of *in vitro* bioactivity over the unbound plasma concentrations to do so. Using the unbound plasma concentrations retrieves known associations with greater strength of association (median Likelihood Ratio) and the associations are more precise (median PPV). However, this is at the cost of a lower recall, as only 33 versus 80 previously reported target-AE associations are retrieved, and a lower detection rate of AE-associated drugs (fraction of AE-associated drugs that are active). Boxplots show the interquartile range (IQR) including the median and whiskers extend to 1.5 times the IQR.

### Quantification of associations between the activity of drugs on proteins and the occurrence of AEs

We next evaluated the bioactivities at protein targets as predictors of AEs by analysing to what extent statistically significant target-AE associations provide information about the AE (Fig. 3A). Therefore, we focus on the PPV, the fraction of active drugs that are associated with the AE, because this relates the *presence* of bioactivity to the *presence* AEs. The median PPV for significant target-AE associations is 0.38 with a STD of 0.2 for SIDER, while FAERS associations have a median PPV of 0.23 (STD = 0.1), meaning that across significant target-AE associations ~23-38% of drugs active at the target are associated with an *in vivo* AE (Fig. 3A). The lower PPVs observed for FAERS compared to SIDER mean that bioactivities have higher false positive rates for AEs in FAERS, and FAERS associations also have lower LRs with a median 11.8 versus 16.4 for SIDER (Fig. 3A). These observations could be due to FAERS being noisier and containing more uncertain AEs, making it more difficult to identify target-AE associations in the dataset.

**Fig. 3.**
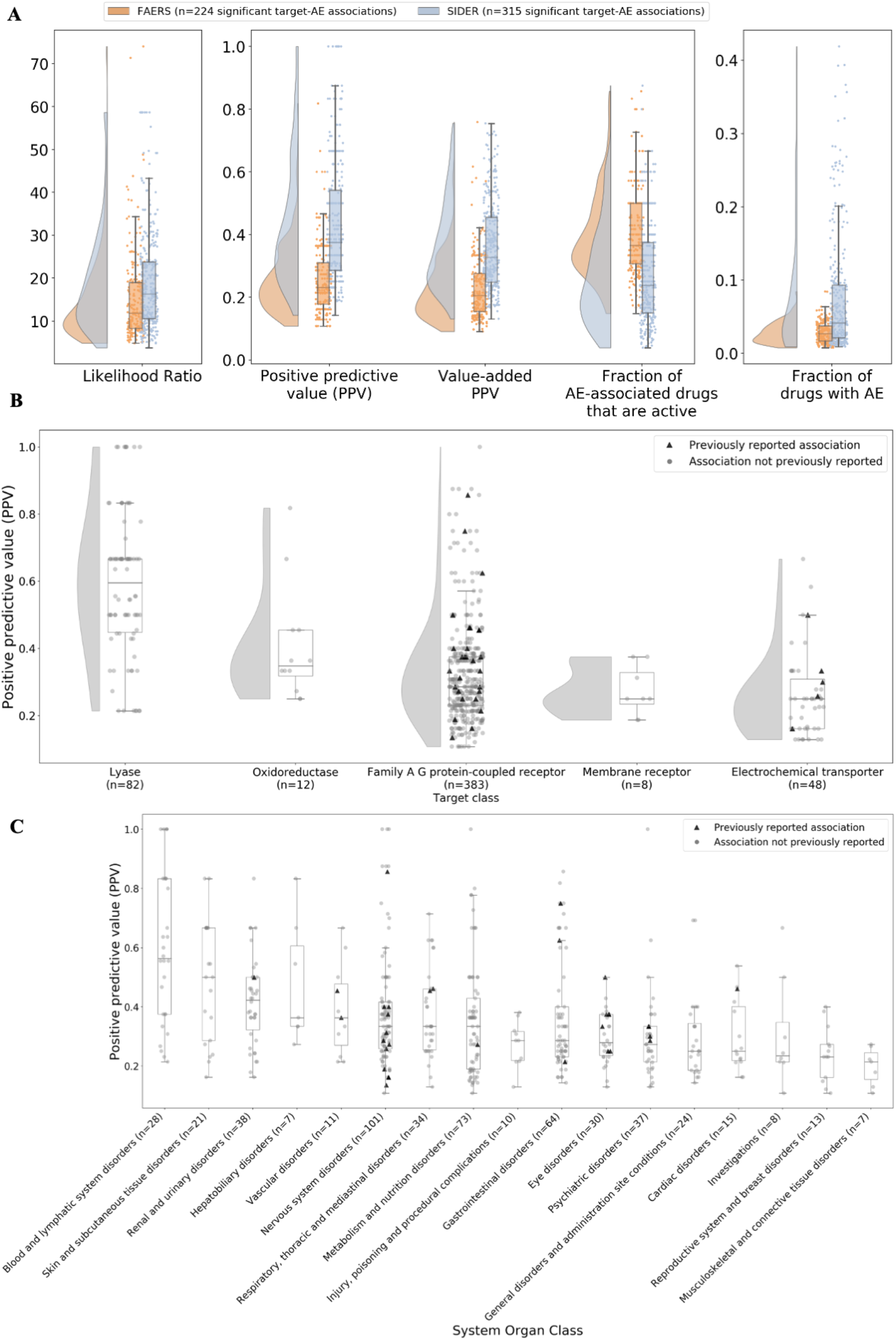
Quantification of statistically significant target-AE associations. (**A**) Association metrics of significant associations in the FAERS and SIDER datasets. Infinite Likelihood Ratios (LRs) have been set to the highest observed value in the respective dataset. Associations in SIDER have higher LRs, with a median of 16.4 compared to 11.8 for FAERS. Similarly, the PPVs are also higher in SIDER, with a median of 0.38 compared to 0.23 for FAERS. (**B**) PPVs of target-AE associations by protein class of the target, for classes with more than 5 target-AE associations. Classes correspond to the second level of the ChEMBL target hierarchy except ‘Membrane receptors’, which is the highest level since those targets were not further classified. The highest PPVs occur in the lyase and family A GPCR classes, which also contain the highest number of target-AE associations. (**C)**PPVs of target-AE associations across MedDRA System Organ Classes of the AEs. The highest median PPV of 0.56 occurs in ‘blood and lymphatic system disorders’. PPVs above 0.8 only occur in the SOCs with the most target-AE associations, such as nervous system and gastrointestinal disorders. Across (**A**), (**B**), and (**C**), boxplots show the IQR including the median and whiskers extend to 1.5 times the IQR.

We also calculated the value-added PPV *(23)*, indicating the additional information provided by the protein target activity over the prevalence to predict AEs. This is because higher PPVs are generally seen with higher prevalence, referring in our study to the fraction of drugs with bioactivity data that are associated with a certain AE *(24)*. The value-added PPVs follow a similar distribution to the PPVs, except the extremes being adjusted such as PPVs close to 1.0 (Fig. 3A). This shows that the target bioactivities provide additional information and that the PPV values are not primarily driven by the prevalence itself. The prevalence (fraction of drugs associated with AE) is plotted for reference, showing that for statistically significant associations the prevalence is generally less than 5% (Fig. 3A).

### Positive predictive values by targets class

We next investigated how PPVs for the target-AE associations differ by target class, shown in Fig. 3B. The highest PPVs (up to 1.0) are observed in the lyase and family A GPCRs, the target classes containing the highest number of associations. Oxidoreductases and electrochemical transporters generally have lower PPVs (up to 0.67), and there are also fewer target-AEs associations in these classes. In addition, a small number of membrane receptors – those not further classified in the ChEMBL hierarchy – also have lower PPVs (up to 0.38) (Fig. 3B). The target-AE pairs that were reported on published safety target panels *(2, 6, 7)* and retrieved in our study are also shown in Fig. 3B, showing that some previously reported associations have low PPVs, such as the relationship between the muscarinic M3 receptor (a family A GPCR) and miosis, which has a PPV of 0.25 (Fig. 3B). This shows that some target-AE associations that have been previously reported were retrieved as significant in our study but have low PPVs, possibly due to AEs being known but not often reported. In other cases the mapping categories used for annotating associations as previously reported, i.e. the MedDRA HLTs, are very broad such as ‘neurological signs and symptoms’ which results in precision being lost. At the same time, the results show that our study identifies many novel target-AE associations which might include mechanistic target-AE links.

### Positive predictive values by System Organ Class (SOC)

We next examined the distribution of PPVs for target-AE associations across MedDRA SOCs of the AEs (Fig. 3C). The range of PPVs within most SOCs is wide, with the highest median PPV of 0.56 occurring for ‘blood and lymphatic system disorders’, and the lowest median PPV of 0.21 for ‘musculoskeletal and connective tissue disorders’ (Fig. 3C). PPVs above 0.8 only occur in the large SOCs such as ‘nervous system disorders’ and ‘gastrointestinal disorders’, which are also the SOCs with the highest number of AEs in the underlying datasets (Fig. S 2). Otherwise, the association metrics do not follow an easily interpretable pattern by SOC (Fig. 3C).

### Trade-off between PPVs and detection of AE-associated drugs

We next analysed the relationship between the value-added PPV and the fraction of AE-associated drugs that are active at the target, which is the fraction of AE-associated drugs that would be detected by bioactivity at the target if used as single predictor of the AE. There is a clear inverse relationship between the two variables (Fig. 4). For example, the value-added PPV of the muscarinic M2 receptor activity for tremor is 0.66, but the fraction of AE-associated drugs of around 0.05 indicates that only 5% of drugs associated with this AE are active at the receptor. Conversely, activity at dopamine D2 receptor would detect 57% of drugs associated with hyperprolactinaemia, but has a false positive rate of 87% (PPV=0.13) (Fig. 4). Specifically, there are few target-AE associations with a fraction of AE-associated drugs that are active above 0.5, and at the same time high PPVs (Fig. 4), which would correspond to a simultaneous high fraction of drugs detected and a low false positive rate, and be most useful in practice. Thus, we can conclude that no single bioactivity is a strong indicator of clinical and post-marketing AEs in the datasets studied. There are many possible reasons for this, e.g. a low fraction of AE-associated drugs would be detected per target if multiple mechanisms involving different targets lead to the same AE, which we will explore further on in the Results. On the other hand, low PPVs can result from pharmacokinetic behaviour of drugs such as lack of blood-brain barrier crossing, leading to certain AEs not being observed in practice.

**Fig. 4.**
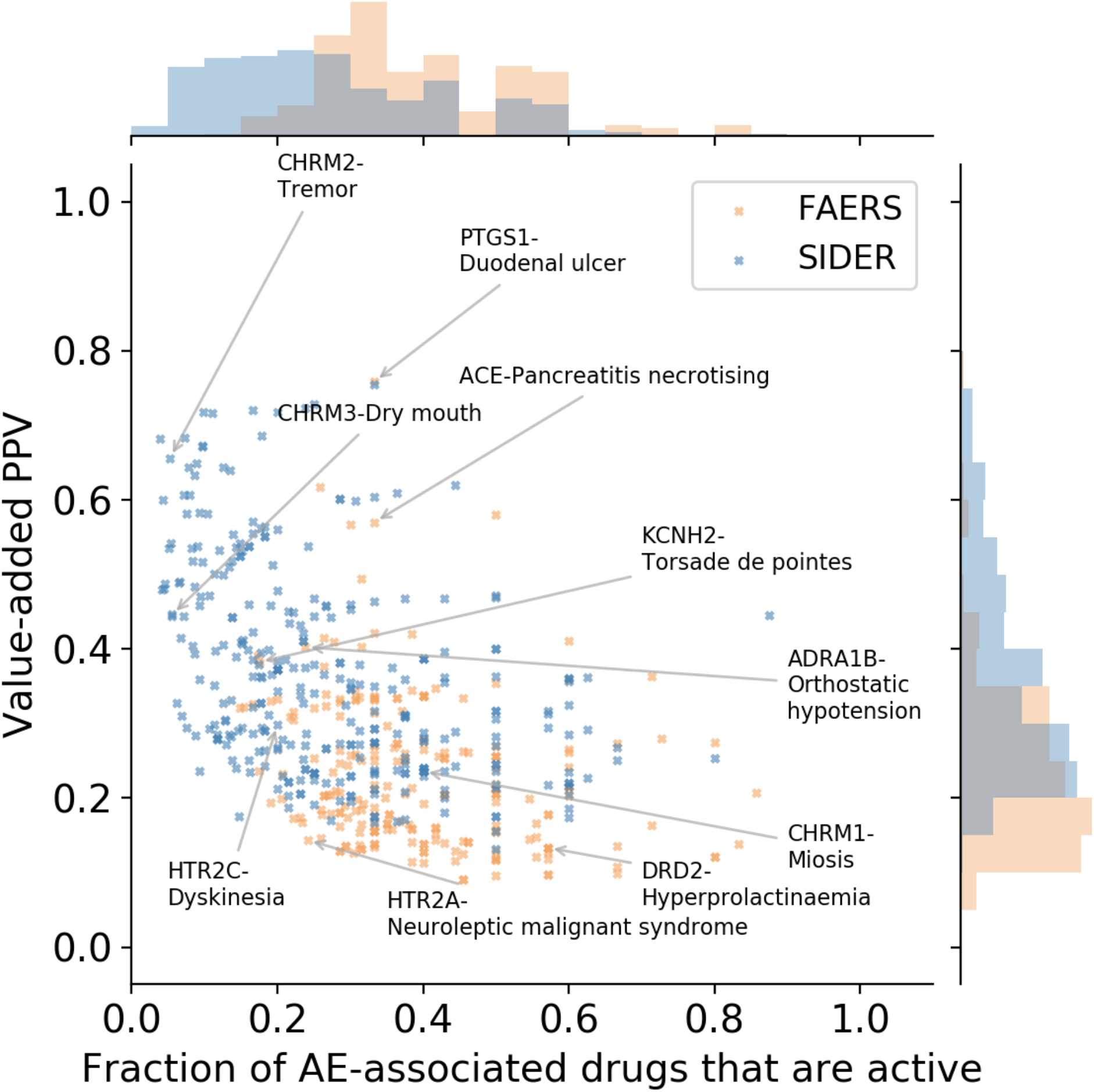
Trade-off between the value-added PPV of significant target-AE associations and the fraction of AE-associated drugs that are active at the target – defined by the ratio of measured or predicted *in vitro* bioactivity and the unbound plasma concentration - and would therefore be ‘detected’ by the target bioactivity. Target-AE pairs with a high value-added PPV tend to have low fractions of AE-associated drugs being active, meaning only a small share of all drugs associated with the AE would be detected by bioactivity at the target. Alternatively, associations with high fractions of AE-associated drugs that are active tend to have low value-added PPVs, indicating a high false positive rate for that target-AE pair. CHRM2: Muscarinic acetylcholine receptor M2, PTGS1: Cyclooxygenase-1, CHRM3: Muscarinic acetylcholine receptor M3, ACE: Angiotensin-converting enzyme, KCNH2: hERG, ADRA1B: α-1b adrenergic receptor, CHRM1: Muscarinic acetylcholine receptor M1, DRD2: Dopamine D2 receptor, HTR2A: Serotonin 2a (5-HT2a) receptor, HTR2C: Serotonin 2c (5-HT2c) receptor.

### Global analysis of observed target-AE associations, with a focus on established and novel safety targets

We next compared the significant targets in our study to previously reported safety targets. In total, our study considered 104 targets out of which 45 were found to have at least one statistically significant association to an AE (Fig. 5). 30 out of these 45 targets are already included on secondary pharmacology panels that we compiled from previous literature *(2, 6, 7)*, so the majority of significant targets in our study are established safety targets (Fig. 5). At the same time, out of the 91 safety targets from literature, for only 40 targets was data available in our study, highlighting the lack of publicly available experimental data for previously reported safety targets (Fig. 5). The remaining 15 targets with significant associations in our study, among which eight are members of the carbonic anhydrase (CA) family, are not included on current panels. Apart from the CAs and one other novel target, namely microtubule-associated protein tau, all other six novel targets are additional family members of proteins already included on current panels such as serotonin, dopamine, and adrenergic receptors, e.g. dopamine D3 receptor is not currently included on panels but dopamine D1 is. The following sections will discuss individual associations in more detail.

**Fig. 5.**
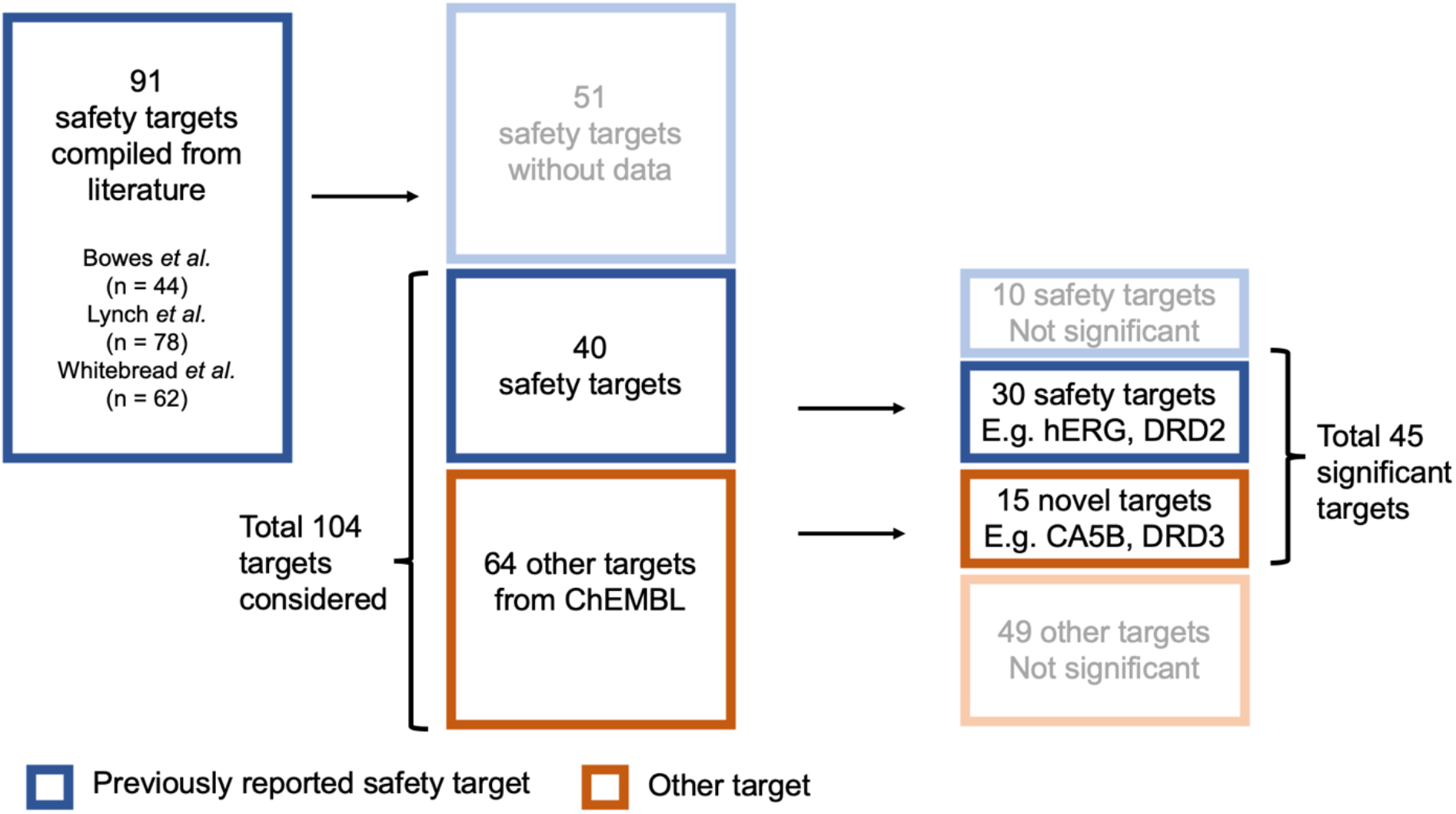
Overview of targets in the study, showing the number of targets considered, those found significantly associated with AEs, and those previously reported on safety target panels. For 51 out of 91 safety targets from literature, no experimental data was available in the current study. Of the 45 targets with significant associations to AEs, 30 are listed on current safety target panels whereas 15 are not and thus potentially novel. CA5B = carbonic anhydrase 5B, DRD2 = dopamine D2 receptor, DRD3 = dopamine D3 receptor.

### Target-AE associations with the highest predictive values

To examine the individual target-AE associations with the highest PPVs, Table 1 shows the significant associations with the highest value-added PPVs in SIDER and FAERS. The table focuses on events within the SOCs of ‘nervous system disorders’, ‘vascular disorders’, ‘cardiac disorders’, and ‘respiratory, thoracic and mediastinal disorders’, because their link to the function of vital organs makes these a high priority in drug safety *(2, 12)*, and ‘hepatobiliary disorders’, which is a leading cause of clinical attrition *(25)*. We will discuss the most highly ranked associations here, but the full list of associations can be found in Data File S 3 (FAERS) and Data File S 4 (SIDER).

**Table 1.**
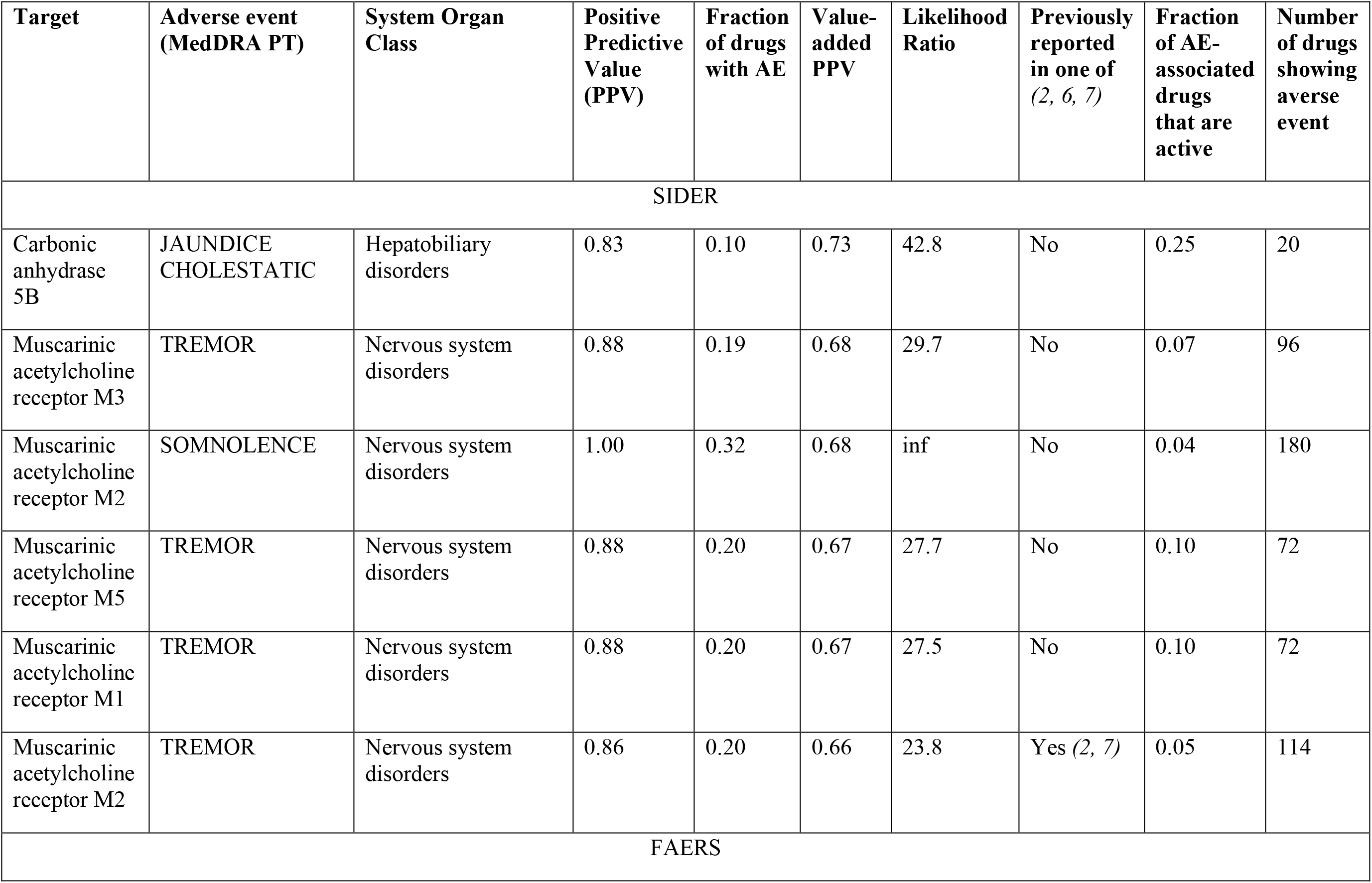
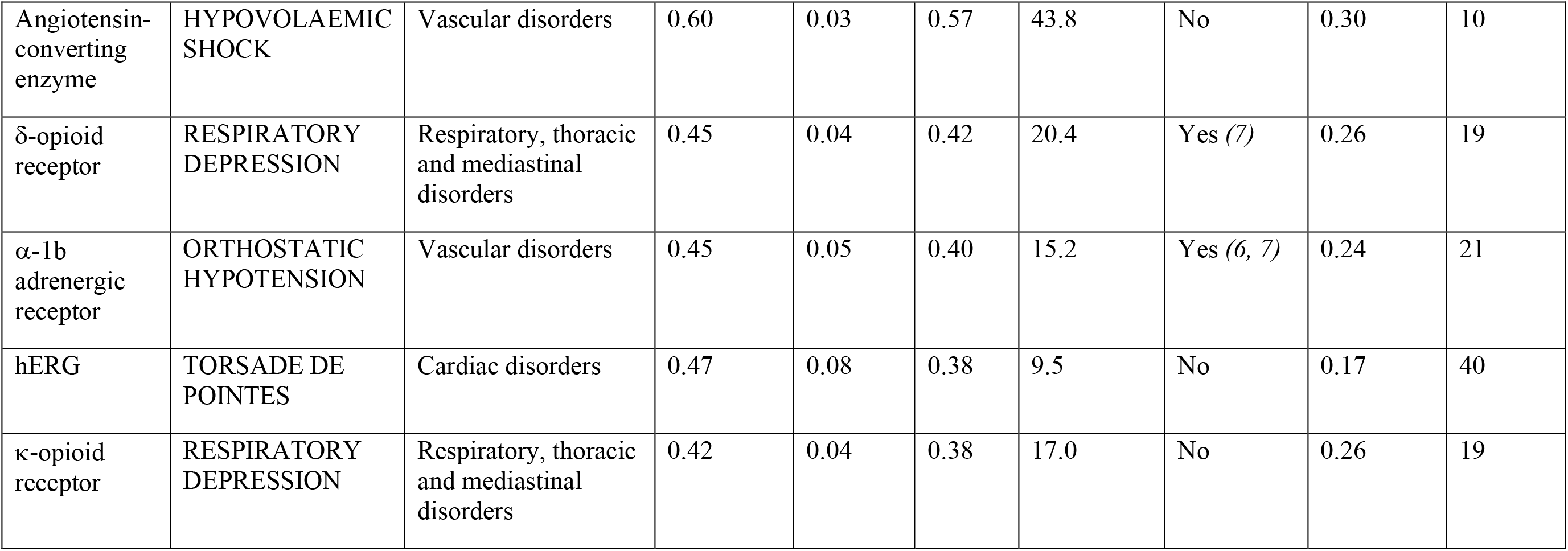
Most highly ranked associations between activity at a protein and observed AEs, sorted by their value-added PPVs, based on the SIDER and FAERS datasets.

Based on SIDER, the most predictive target-AE association is between CA5B and cholestatic jaundice, a liver disorder, with a PPV of 0.83 (Table 1). The CA family of enzymes are involved in acid-base balance and CA inhibitors are used clinically in a range of conditions including ocular disorders, oedema, and seizures, and they are often not entirely selective across CAs *(26–29)*. The association between CA5B and cholestatic jaundice is plausible due to their high tissue expression in the mitochondria *(30)*, which is relevant to liver toxicity *(31)*. We are not aware of a direct link between CAs and cholestatic jaundice being previously reported, although CA4 and CA12 were identified as protein interactors of targets associated with cholestatic jaundice the study by Duran-Frigola and Aloy, suggesting the existence of additional indirect links via the protein-protein interaction network *(11)*.

The next most predictive associations concern various muscarinic acetylcholine receptors. In our study, the muscarinic receptor M2 is associated with somnolence (PPV=1.0, Table 1), which is in line with acetylcholine being important for wakefulness and the expected effects of M2 antagonism *(32)*. In practice, somnolence is a common side effect of muscarinic acetylcholine M2 and M3 receptor antagonists such as oxybutynin and tolterodine *(33–35)*. Next, we identified that the muscarinic acetylcholine receptors M3, M5, M1 and M2 are each associated with tremor with similar PPVs between 0.86-0.88 in our results (Table 1). The link between the muscarinic acetylcholine receptors M2 and tremor has been previously reported *(2)*, and the links of multiple receptors in our study could either be due to multiple targets being biologically related to the effect, or compound promiscuity since the muscarinic receptors share between half and all their active ligands with each other in our dataset (Fig. S 5A). In contrast to the CAs, the muscarinic M1 and M2 receptors are included on all published safety panels we considered and M5 on the Lynch panel only *(2, 6, 7)*. Hence, we conclude that the most predictive target-AE associations based on SIDER include a novel link for CA5B, while other associations are previously reported and in line with current mechanistic knowledge and previous literature. However, several links may be driven by compound promiscuity, which cannot be distinguished based on the current statistical analysis.

For the analysis of the FAERS database, the association with the highest value-added PPV is between the angiotensin-converting enzyme (ACE), a target included on the Lynch panel only *(7)*, and hypovolemic shock, which is circulatory failure due to fluid loss (PPV=0.60, Table 1). This is consistent with the fact that ACE inhibitors interfere with the renin-angiotensin system that normally protects against hypovolemia *(36)*. There are case reports in literature that have attributed hypovolemic shock to ACE inhibitors *(37)*.

The known associations between the δ-opioid receptor and respiratory depression *(7)* and between the α-1b adrenergic receptor and orthostatic hypotension *(7)* were successfully retrieved in the current analysis with PPVs of 0.45 (Table 1). The δ-opioid receptor is included on all considered safety panels and the α-1b adrenergic receptor on the Whitebread and Lynch panels *(6, 7)*. Similarly, the association between hERG, one of most studied and screened safety targets *(2, 38, 39)*, and torsade de pointes (TdP) is retrieved with a PPV of 0.47 (Table 1). A previous study by Pollard et al. found a PPV of 1.0 for the relationship between hERG and QT interval prolongation when the margin between the *in vitro* hERG IC50 and C_max_ was less than 10-fold *(38)*. The difference between this PPV and the one in our study is explained by QT interval prolongation being a risk factor for TdP but TdP being the ultimate ventricular arrhythmia *(2, 38, 40)*, since a considerable number of drugs cause QT interval prolongation but are not torsadogenic *(39)*. Only QT prolongation is listed among the compiled previously reported associations *(2, 6, 7)* and it has a different MedDRA HLT (investigations) as TdP (ventricular arrhythmias and cardiac arrest), resulting in the association between hERG-TdP in our study not being annotated as previously reported (Table 1). This highlights the challenges in using medical terminologies on a large scale, since descriptions of biological effects may not directly match all related AE terms *(41)*. The next association in our results is between respiratory depression and the κ-opioid receptor (PPV=0.42), which is included on the Bowes and Lynch panels *(2, 7)*.

While activation of μ-opioid and δ-opioid receptors causes respiratory depression *(7, 42)*, the κ-opioid receptor is believed to lack this effect *(42)*. However, the presence of this association in our study can be explained by compound promiscuity, given the high overlap of shared ligands between the opioid receptors (Fig. S 5B). In conclusion, among the most predictive associations in FAERS there are of associations supported by previous literature and associations due to compound promiscuity.

### Novel associations

We next analysed the significant target-AE associations for targets other than those already listed on the previous screening panels considered *(2, 6, 7)*, to identify whether any of these novel targets could provide additional information to predict AEs within the high-priority SOCs listed earlier. Therefore, we will first considered novel targets without family members on current panels *(2, 6, 7)*. These foremost are the CAs which are associated to a range of AEs in addition to the link between CA5B-cholestatic jaundice discussed earlier. All the effects associated with CAs are unique to this target family, meaning no other targets are associated to the same effects in our dataset. Based on SIDER, CA5A (PPV=0.56), CA12 (PPV=0.45) and CA9 (PPV=0.26) are also associated to cholestatic jaundice, of which CA5A is most plausible because of its high liver expression *(43)*. Furthermore, CA9 (PPV=0.31) and CA12 (PPV=0.24) are associated with hepatic necrosis, adding further evidence to the link between CAs and liver effects. CA5A and CA5B are both associated with paraesthesia (PPV=1.0), which is listed as a side effect of CA inhibitors and has been suggested to be caused by CA activity *(27, 44)*. Based on the FAERS analysis, CA4 is associated with hyperammonaemic encephalopathy (PPV=0.38), which is consistent with mechanistic knowledge of CA inhibitors on ammonia balance *(27)*. Lastly, CA2 is associated with simple partial seizures, and CA5B with pulmonary oedema, but both are most likely examples of indication bias in FAERS. Overall, since none of the existing panels include any members of the CA family and all the associated AEs are unique to this family, our results suggest that CAs might be able to extend the coverage of future safety target panels.

The other target without family members on existing panels is microtubule-associated protein tau, which is associated to liver injury with a PPV=0.28 based on FAERS. This protein is associated with neurotoxicity and is currently not the therapeutic target of any approved drug. Normal phosphorylation of microtubule-associated protein tau is disturbed by the microcystin group of bacterial toxins, which are also associated with hepatotoxicity *(45)*, providing some support for our observation. Thus, this link could also be a novel association of interest for safety screening.

Now turning to links between novel targets that are family members of currently screened targets, we found that in nearly all cases both novel and established targets are associated with the same AE. For example, the 5-HT6 receptor, not currently included on any of the considered panels, is associated with tremor (PPV= 0.71), but the muscarinic M3, M5 and M1 receptors are associated to the same effect with a higher PPVs of 0.88 (Table 1). The only exception is α-1d adrenergic receptor’s association to loss of consciousness with PPV= 0.54 based on SIDER, an AE to which no other target is associated in our study. In all the other cases based on SIDER, the novel target had lower or the same PPVs and LRs as a currently screened target. Similarly, based on FAERS, novel targets had lower or comparable PPVs in all cases. This shows that different targets can provide similar levels of information about the same AE and might be redundant in the context of safety target screening if there is a large overlap in active drugs across targets. The extent of promiscuity will determine whether novel targets can provide additional information to improve the detection of AE-associated drugs, which we will discuss next.

### Considering activity against multiple protein targets for the same AE improves the detection of AE-associated drugs

To analyse the value of considering activity against multiple proteins associated with the same AE – using a logical ‘or’ – and find targets that provide non-redundant information, we examined combinations of targets that can increase the detection of AE-associated drugs. In 38% (FAERS) and 45% (SIDER) of AEs, we found that considering activity at one of multiple targets improves the detection of AE-associated drugs. Generally, considering two or three targets associated with the AE leads to a median improvement of 20% (FAERS) and 33% (SIDER) in the detection of AE-associated drugs (Fig. 6A). This is at a cost of worsening PPV by a median 16% (FAERS) and 21% (SIDER) (Fig. 6A). The improvements are due to each of the targets identifying different AE-associated drugs. The combination with the greatest improvement in detection compared to the single target is the combination of the β-1 adrenergic receptor and CA5B in relation to orthostatic hypotension in case of the SIDER dataset (Fig. 6B). Both targets individually detect 12% of AE-associated drugs, but this is increased to 25% when considering activity at either one of the targets (Fig. 6B). It is not surprising that a CA is involved in the best performing target set, since active drugs at the CAs generally overlap little with those against other targets in the study (Fig. S 5).

**Fig. 6.**
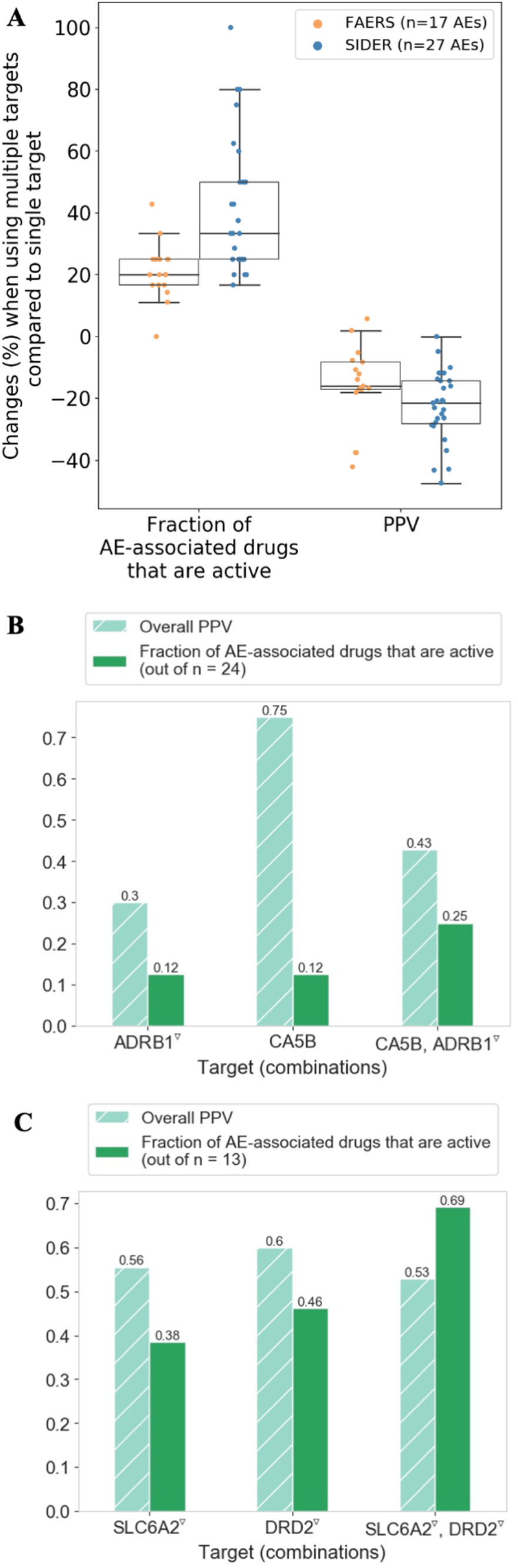
(**A**) Changes in the PPV and fraction of AE-associated drugs active at the target (‘detected’) when comparing combinations of targets – i.e. activity at either target – for a given AE to the individual targets. Considering activity at multiple targets improves the detection of AE-associated drugs considerably, at the cost of decreasing PPV. Thus, considering activity against multiple targets can help anticipate AEs, while detailed mechanistic investigations are beyond the scope of our study. (**B**) PPV and fraction of drugs associated with orthostatic hypotension that are active at the β-1 adrenergic receptor (ADRB1) and carbonic anhydrase 5B (CA5B), as individual targets and in combination, based on 143 drugs that were measured consistently at both targets. (**C**) PPV and fraction of drugs associated with Neuroleptic Malignant Syndrome that are active at the dopamine D2 receptor (DRD2) and norepinephrine transporter (SLC6A4), as individual targets and in combination, based on 320 drugs that were measured at both targets. In (**B**) and (**C**), the target combinations detect the highest number of AE-associated drugs.

The AE with the highest overall percentage of AE-associated drugs detected by a combination of proteins is Neuroleptic Malignant Syndrome in the SIDER dataset (Fig. 6C); considering activity at either the dopamine D2 receptor or the norepinephrine transporter detects 69% of AE-associated drugs with an overall PPV of 0.32. These targets are consistent with currently known mechanisms behind Neuroleptic Malignant Syndrome *(46)*.

In conclusion, in about 40% of AEs in our study, considering activity at each of a sets of targets associated with the same event can improve the detection of AE-associated drugs by a median one-fifth to one-third, with generally lower decreases in PPV, and an analysis of these combinations can identify targets that provide orthogonal, as opposed to redundant information.

### Fraction of AEs associated with protein activity and hence potentially detectable from safety pharmacology screens

We next investigated what fraction of unique AEs in the dataset have at least one significantly associated target in order to estimate what fraction of such events may be detectable from protein-based safety pharmacology screens. We found that 8.5% of unique AEs in the SIDER dataset and 2.9% in the FAERS dataset have one or more significantly associated target. These low percentages are partially due to the use of unbound plasma concentrations, which restricted the total amount of data available for analysis, since the percentages using the constant cut-off are 44.1% (SIDER) and 19.6 (FAERS). The fact that the percentage is lower for FAERS could be related to the presence of biases and noisier nature of FAERS reporting *(16)*. FAERS also contains a greater diversity of AEs related to a similar number of drugs compared to SIDER (Fig. S 2), and target-AE associations are less likely to be detected when only a few drugs are associated such as to rare events *(10)*.

### System Organ Class distribution of AEs associated with targets

To examine differences in how AEs belonging to different SOCs are associated with targets, we compared the fraction of unique AEs in the underlying dataset of drug-AE associations – including AEs that could not be statistically related to targets – to the AEs that were present among the statistically significant target-AE associations (Fig. 7). This shows that AEs in some SOCs are more frequently associated with targets than AEs from other classes. For example, the largest percentage difference is observed for AEs in the ‘metabolism and nutrition disorders’ class, which comprise 3.9% (SIDER) and 2.7% (FAERS) of unique AEs in the underlying datasets, but 9.4% (SIDER) and 10.3% (FAERS) of AEs statistically associated with targets (Fig. 7). The next largest overrepresented classes are gastrointestinal, nervous system, and psychiatric disorders in FAERS, and nervous system, ‘blood and lymphatic system’, and ‘respiratory, thoracic and mediastinal’ disorders in SIDER. The enrichment of ‘metabolism and nutrition disorders’ and gastrointestinal (GI) AEs suggest these can more often be related to pharmacological actions, as suggested by familiar examples being retrieved in our study, such as cyclooxygenase-1 and gastric ulceration *(2)*, and muscarinic acetylcholine receptor M3-mediated dry mouth and constipation *(2)*. GI disorders also have one of the largest number of drugs associated with them (Fig. S 2), thus forming a larger dataset for statistical discovery. AEs in some of the above enriched SOCs also ranked highly for correspondence to target phenotypes in the study by Deaton et al. *(12)*, such as platelet disorders and nonhaemolytic anaemias (blood and lymphatic system), and glucose metabolism disorders (metabolism and nutrition disorders).

**Fig. 7.**
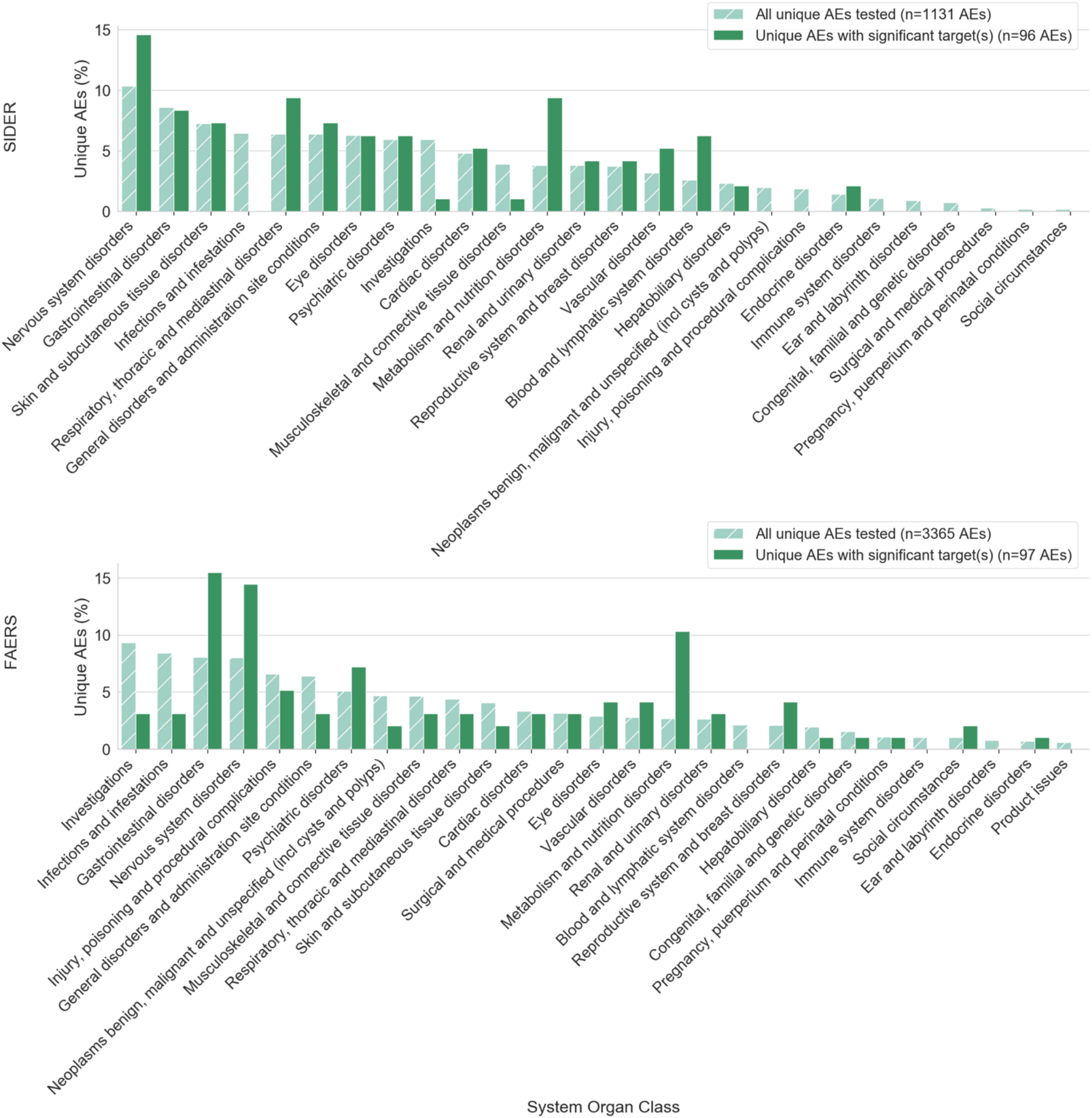
Relating AEs to targets by SOC, comparing the AEs in the underlying drug-AE datasets for (A) SIDER and (B) FAERS to those AEs that are statistically associated with targets in our study. AEs in some classes (e.g. ‘metabolism and nutrition disorders’) are more often associated with targets, whereas AEs in other classes are present in the dataset but rarely associated with targets (e.g. ‘infections and infestations’).

Overrepresentation of nervous system, psychiatric, and respiratory disorders noted above could be related to the prominent presence of GPCRs in the dataset, which frequently target these organ systems and neurotransmission generally *(2, 47)* In contrast, the largest underrepresented class is ‘investigations’, which makes up 5.9% (SIDER) and 9.3% (FAERS) of AEs in the underlying dataset, but only 1.0% (SIDER) and 3.1% (FAERS) of the set related to targets (Fig. 7). AEs in this category are sometimes relatively unspecific such as ‘blood test abnormal’, which might be a reason for the lack of associations to targets. Similarly, the next most underrepresented classes are ‘musculoskeletal and connective tissue disorders’ for SIDER and ‘infections and infestations’ in FAERS. Many mechanisms by which drugs can increase susceptibility to infections, such as immunosuppression due to cytotoxicity *(48)*, or disruption of the gut microbiota *(49)*, are not covered by *in vitro* pharmacology, explaining their underrepresentation. Overall, we conclude that target-AE associations are not uniformly distributed across SOCs, with classes such as ‘metabolism and nutrition disorders’ and nervous system disorders being most frequently related to pharmacological targets.

### Protein activities are frequently associated with different AEs in FAERS and SIDER

We next investigated to what extent FAERS and SIDER are complementary for identifying target-AE associations, and to this end we calculated the overlap in AEs associated with the same targets in either dataset (Table S 1; see Data File S 3 and Data File S 4 for full list of associations). The highest overlap in unique PTs between the datasets is 7% for the dopamine D2 receptor (Table S 1). Considering the HLT increases the overlap to some extent, but apart from hERG, which is associated with ‘ventricular arrhythmias and cardiac arrest’ in both datasets, leading to 100% overlap in HLTs, the next highest overlap is still low at only 10% for the dopamine D2 receptor (Table S 1). This means that in our study, the FAERS and SIDER datasets are associated with nearly completely disjoint sets of AEs for the same targets, thus being highly complementary. The complementarity could reflect different AEs being reported in clinical trials versus post-marketing phases, for example due to more detailed patient observation in clinical trials, or due to or differences in short and long-term drug effects. The latter is also supported by the observation that FAERS is the only dataset providing target-AE associations in some SOCs, such as neoplasms and ‘pregnancy, puerperium and perinatal conditions’ (Fig. 7), which could be related to long-term use and more diverse populations exposed in the post-marketing phase.

Other reasons for differences in reported AEs are that AEs already listed on a drug label are less likely to get reported to post-marketing systems *(50)*, and the inclusion of more uncertain reports in FAERS due to lack of causal evidence and submission by patients as opposed to healthcare professionals *(15, 16)*. Thus, we conclude that FAERS and SIDER provide different target-AE associations, and each could provide added value for detecting target-AE associations.

## Discussion

In this work we identified and quantified associations between drugs’ pharmacological activities and AEs observed in clinical trials and during post-marketing surveillance. Compared to previous studies, our study for the first time takes into account unbound drug plasma concentrations on a systematic scale.

We found that taking into account drug plasma concentrations reduced the size of the dataset by about half, but increased the PPV and LR of associations reported in previous literature at the cost of lower recall. This suggests that integration with plasma concentrations is more precise but recall is limited by data availability.

Many of the most predictive associations in our study are supported by previous literature, but some associations appeared due to compound promiscuity, which remains a source of potential false positive associations in studies of statistical nature. Thus, regarding any statistical associations derived in our study, further research would be needed to confirm mechanistic links. Key novel findings such as the association of CA5A and CA5B with liver effects would be suggested for further investigation and future inclusion in safety target panels.

No single target-AE association had a PPV above 0.5 while at the same time identifying more than 50% of drugs associated with a given AE, showing that single *in vitro* bioactivities do not perfectly indicate *in vivo* effects. However, we found that considering activity at multiple targets associated with the same event can improve the detection of AE-associated drugs in about 40% of cases. The lack of one-to-one translation between *in vitro* and *in vivo* effects in our study corroborates previous findings, for example those that observed a lack of correlations between large-scale *in vitro* bioactivities and toxicity observed in animal studies *(51, 52)*, and those reporting high false positive rates associated with early screening for targets such as hERG and BSEP *(38, 39, 53)*. Reasons for the remaining challenge to translate *in vitro* to *in vivo* effects include differences between plasma and tissue concentrations *(13)*, varying protein expression across tissues, and interactions between targets and pathways, such as transporter and ion channels off-setting each other’s effects *(39, 53)*. Our finding that more AE-associated drugs can be identified by using combinations of targets as opposed to single targets is consistent with the knowledge that AEs can be caused by multiple mechanisms involving different targets. Although we only explored combinations of targets with the ‘or’ operator, *simultaneous* modulation of targets – ‘and’ operator – may be involved in some AEs, which would not have been discovered in our study.

Regarding factors that influence the association between bioactivities and AEs, we found no strong patterns across target class or SOC of the AE in terms of PPVs, but we did find that AEs in some classes were more frequently associated to targets, in particular ‘metabolism and nutrition disorders’, GI disorders, and nervous system disorders, suggesting that based our data AEs in these SOCs are more frequently related to pharmacological targets and hence more likely detectable from protein-based screening. However, in total, only a small fraction of all unique AEs in FAERS (2.9%) and SIDER (8.5%), could be statistically related to at least one target.

This is lower than the 75% of reported AEs being predictable from pharmacology *(5, 54)*, but these latter estimates are based on the frequency of the AE in the population, whereas our numbers look at unique AEs. The low percentages are partly due to the smaller subset of data with plasma concentrations, since the percentage is up to 44% when using an absolute pChEMBL cut-off ≥ 6 (1 μM). That figure is similar to the 51% of AEs being related to targets in the study by Bork et al. *(10)*. However, as a result of the current study we would conclude that those numbers are a result of trading PPV for recall, the desirability of which depends on the particular situation.

There are several limitations in our study that may have resulted in the low PPVs and fractions of AE-associated drugs detected. Firstly, since plasma concentrations were only available for a subset of the data, this likely reduced the power of our study. It is also well known that bioactivity datasets are sparse and incomplete, further limiting the power and recall *(21)*. Our analysis did not take into account functional effects, such as agonism or antagonism, as this information is not consistently available from the databases considered here *(2, 21)*. This may have resulted in the masking of associations only associated with certain functional effects or modes of action. Similarly, the effects previously reported in e.g. Bowes et al. generally do not list the dose at which effects are expected, so effects only seen at high dose may be missed *(2)*, and this may also explain some of the discrepancies between our results and previous studies.

Furthermore, our analysis is based on marketed drugs which have already undergone safety screening, so the termination of problematic candidates early on has biased the data available to us in a way that strong associations with safety targets may not be apparent. There are uncertainties about the drug-AE links on which this work is based, since in FAERS associations can be purely statistical and not causal *(16)*. Lastly, limitations of the drug plasma concentrations used in our study include that they are a mixture of ‘normal’ therapeutic concentrations and C_max_, that they are not patient-specific, and do not distinguish between different drug indications that may require different doses of the same drug.

Our study showed that FAERS and SIDER are complementary with respect to signals detected, because the same targets were associated with different AEs in either dataset. A previous study considered solely AEs reported in both FAERS and SIDER *(12)*, but our results suggest this approach would potentially result in a large part of the AE space not being considered.

Our results are relevant to current developments in predictive toxicology, especially the focus on applying machine learning to predict target bioactivities for lists of toxicity-related targets or assays *(55–57)*. The better we are able to associate protein activities with AEs, and the better we understand the pharmacokinetics of drugs, the better we will be able to translate *in vitro* into *in vivo* effects.

## Materials and Methods

### FAERS

The FAERS AEOLUS database, containing 3,526 drugs mapped to the RxNorm drug vocabulary and 17,710 unique AE terms, was installed as a MySQL database *(58)*. Drugs were mapped to ChEMBL parent compounds, which group different salt forms of the same drug and link to a unique molecular parent structure, using the mapping between RxNorm concepts and DrugBank identifiers provided by the RXNCONSO file from the RxNorm vocabulary (RxNorm_full_12032018, downloaded from *(59)*) and UniChem *(60)*, as well as direct matches between RxNorm compound names and synonyms and ChEMBL pref_name, compound_names and synonyms. We were able to map 2,764 drugs to ChEMBL identifiers.

### Propensity score matching

We applied the PSM technique as described by Tatonetti et al. to reduce the effects of confounding factors in the FAERS AEOLUS data (Fig. 1, box 1) *(19)*. The aim is to identify sets of reports that match on underlying patient characteristics but differ in whether the drug of interest is listed, for these reports to be used in disproportionality analysis. Briefly, for each drug, the 200 most strongly correlated concomitant drugs and indications were identified using the Fisher’s exact test and Benjamini-Hochberg correction (corrected p-value <0.05). These covariates are then used in a logistic regression PSM model, of which the dependent variable is whether or not the drug was prescribed to a patient given their characteristics. The model is used to calculate propensity scores for all reports listing the drug (exposed reports) and 100,000 randomly sampled reports not listing the drug (non-exposed/control reports). By using 20 equally spaced propensity score bins, sets of exposed and 10 times as many control reports with matched scores (sampled with replacement) are identified for further analysis.

### Disproportionality analysis on FAERS AEOLUS

Using the matched reports selected using PSM, the rate of AE reporting was calculated in each group of reports, using the MedDRA PT for the AEs from the standard_case_outcome table. The fraction of patient reports listing an AE of interest in the exposed group over the same fraction in the non-exposed reports was calculated to give the PRR (Fig. 1, box 2) *(61)*. The corresponding χ^2^ statistic was calculated using the SciPy chi2_contingency function *(62)*, requiring a minimum of five reports in each cell of the contingency table. Significant drug-AE pairs were identified using an existing pharmacovigilance signal detection threshold of PRR > 2 and χ^2^ > 4 (Fig. 1, box 2) *(63)*. We were able to calculate the PRR for AEs reported for 1,388 mapped drugs.

### Side effects from SIDER

We used the download files provided for SIDER version 4.1 which contain a total of 1,556 drugs *(20)*. Excluding biologicals/peptides, we were able to map 1,219 to ChEMBL identifiers using InChI keys from UniChem *(60)*. To retain the clinical effects only (Fig. 1, box 3), we first retained all MedDRA PTs from the meddra_all_se file, which lists all effects per drug, for drugs that did not have any post-marketing annotations in the meddra_freq file, which only contains information for a subset of drugs. For drugs present in both files, we excluded effects listed as post-marketing, unless a frequency was available for that same effect, since frequencies can generally only be derived from clinical trials. This yielded a set of 1,027 drugs with side effects.

### *Extraction of* in vitro *bioactivity data*

Bioactivities of drugs from the FAERS and SIDER datasets against human single proteins or protein complexes were retrieved from a local MySQL installation of ChEMBL version 24.1 (Fig. 1, box 4). In case of protein complexes, bioactivities at each of the constituent single proteins listed in the target_components table were used. The activity types included were ‘IC50’, ‘EC50’, ‘XC50’, ‘AC50’, ‘Ki’, ‘Kd’ and ‘Potency’ for assay of types ‘B’ (binding) and ‘F’ (functional), with standard_flag = 1 and confidence scores of 7 (protein complexes) and 9 (single proteins), ensuring the highest level of confidence in target assignment *(21)*. Inactive data was retrieved by extracting records containing any of the following in the activity_comment: ‘Not active’, ‘inactive’, ‘No inhibition’ and allowing standard_flag = 0 for these records. The median reported pXC50 was computed per parent drug for further analysis and in case of conflicting active and inactive (based on the activity comment) assignments for the same drug, the active assignment was taken forward.

### Target prediction

The probability of activity at single human proteins at different bioactivity levels was predicted using the target prediction tool PIDGIN version 3 (Fig. 1, box 5) *(22)* as follows: compounds were standardised using the e-Tox standardiser *(64)*, and separate classification models provided predictions for the probability of activity below the thresholds 0.1, 1, 10 and 100 μM. The applicability domain threshold was set to 0.7 and only models with a minimum Precision-Recall Area Under the Curve of 0.7 during cross-validation were included *(65)*. Experimental bioactivity values were used where available, with predictions obtained using the settings described here being added to the compound-target bioactivity matrix. The share of predicted versus measured data points used per target is shown in Data File S 7 (SIDER) and Data File S 8 (FAERS).

### Plasma concentration data

Therapeutic drug plasma concentration data were compiled from a variety of sources (Fig. 1, box 6) and are provided in Data File S 9. We extracted the upper value of the therapeutic plasma concentrations from *(66)*. We also queried ChEMBL 24.1 for human C_max_ data for parent drugs in plasma, blood or serum, using the standard_flag = 1 and excluding rows with data_validity_comment = ‘Outside typical range’. For the plasma concentrations we calculated molar values using the molecular weight of the parent compound from ChEMBL version 24.1. Data for fraction unbound (Fu) and plasma protein binding (PPB) were retrieved from ChEMBL_24_1. Additional Fu, PPB, and unbound plasma concentrations were extracted from several publications *(14, 39, 40, 67)*. For all publications we either used the InChI Key provided or mapped the drug names from the publication to ChEMBL 24.1 drugs via direct matches to ChEMBL pref_name, synonyms, or compound_name. The median Fu values were calculated using Fu and percentage PPB data, and then multiplied by the total plasma concentrations to derive the unbound plasma concentration. Then the median calculated unbound plasma concentration was used for further analysis (Data File S 10).

### *Integration of bioactivity data with drug plasma concentrations to assign expected target engagement* in vivo

The measured and predicted bioactivities of drugs and their unbound plasma concentrations were compared to assign an active or inactive label to each drug-target pair, referred from here onwards as the ‘activity call’. The drug was assigned as active at a given target if the drug plasma concentration exceeded the *in vitro* bioactivity concentration, or was within 1 log unit of the *in vitro* concentration (Fig. 1, box 7). The leeway of 1 log unit is aimed at maximising the potential signal given experimental variability of bioactivity measurements, which may be around half a log unit *(68)*, and variability in pharmacokinetics. If the unbound plasma concentration was more than 1 log unit lower than the measured *in vitro* concentration, the drug was recorded as inactive for a particular target. Additionally, inactive calls were made based on the activity_comment conveying inactivity as described above (Fig. 1, box 7). Assuming that unbound plasma concentrations correspond to concentrations in target tissues, this gives an approximation of whether a target will be modulated sufficiently to obtain an effect.

### Integration of target predictions with drug plasma concentrations

Depending on whether drugs were in the applicability domain of a target prediction model, probabilities between 0 and 1 were available at all or some of the thresholds (0.1, 1, 10 and 100 μM) for each drug-target pair. When predictions were available, a predicted activity probability below 0.4 was considered inactive and above 0.6 active for each threshold. The lowest predicted active threshold was compared to the unbound plasma concentration in the same way as for measured data above (Fig. 1, box 7).

### Identifying significantly associated target-AE associations

For each target, the drugs from the AE datasets with available measured bioactivity data or target predictions were identified (Fig. 1, box 8). Thus, no inactivity was assumed beyond inactive data points and inactive target predictions. The Scikit-learn 0.21.3 *(69)* confusion_matrix function was used, using the *in vitro* bioactivity as predictor of the AE and obtaining the number of true negatives (TN), false positives (FP), false negatives (FN) and true positives (TP) (Fig. 1, box 8) *(70)*. The LR was calculated as LR = TP*(FP + TN)/FP*(TP + FN) *(71)*. The Fisher’s exact p-value was calculated using the SciPy stats 1.3.2 fisher_exact function *(62)*, limited to those associations with at least of five drugs active at a target and five drugs associated with the AE, as in previous studies *(10, 11)*. For all pairs with a positive association (LR > 1), the p-values were corrected for multiple testing with the Benjamini-Hochberg method using the Statsmodels 0.9.0 multipletests function *(72)*. We considered associations significant if they had a corrected p-value ≤ 0.05. The PPV was calculated as PPV = TP/(TP+FP). The value-added PPV was calculated by subtracting the prevalence, i.e. the fraction of drugs measured at a target that is associated with an AE, from the PPV *(23)*.

### Compilation of previously reported safety associations

Protein names and adverse effect descriptions were manually extracted from previous work *(2, 6, 7)*. Protein names were manually mapped to Uniprot identifiers in ChEMBL, based on name. Arrows and terms such as ‘enhances’, ‘induces’, ‘facilitates’, ‘exacerbates’ etc. were reformatted to ‘increased’ and terms such as ‘inhibits’, ‘reduces’, ‘impairs’ etc. to ‘decreased’. The terms were then submitted to the NCBO Bioportal Annotate functionality for annotation with MedDRA terms *(73)*. All results were manually inspected and selected. Additionally, mappings for terms not mapped in this way were identified by manually querying the term in the MedDRA web-based browser and selecting appropriate PTs or HLTs *(74)*. All the mapped associations are provided in Data File S 11 (PT) and Data File S 12 (HLTs). MedDRA HLTs from MedDRA (versions 21.1 or 22.1) were obtained using the Hierarchy Analysis function in the MedDRA web-based browser *(74)*, and associations from FAERS and SIDER in the current study were considered previously reported if the HLT matched a previously reported associations for the same target.

## Acknowledgments

Thanks to Kathryn A. Giblin, Lewis H. Mervin and Will Redfern for helpful discussions. **Funding**: Lhasa Limited, Leeds. **Author contributions**: Conceptualization: AB, TH, IAS, FS, AMA. Data curation: IAS, CHGA, Formal analysis: IAS. Software: IAS, AMA, CHGA. Writing – original draft: IAS Writing – review & editing: AB, TH, FS, IAS, CHGA, AMA **Competing interests**: None. **Data and materials availability**: Code and additional files are available from https://github.com/inessmit/adverse_event_analysis

## Supplementary Materials

**Fig. S1.**
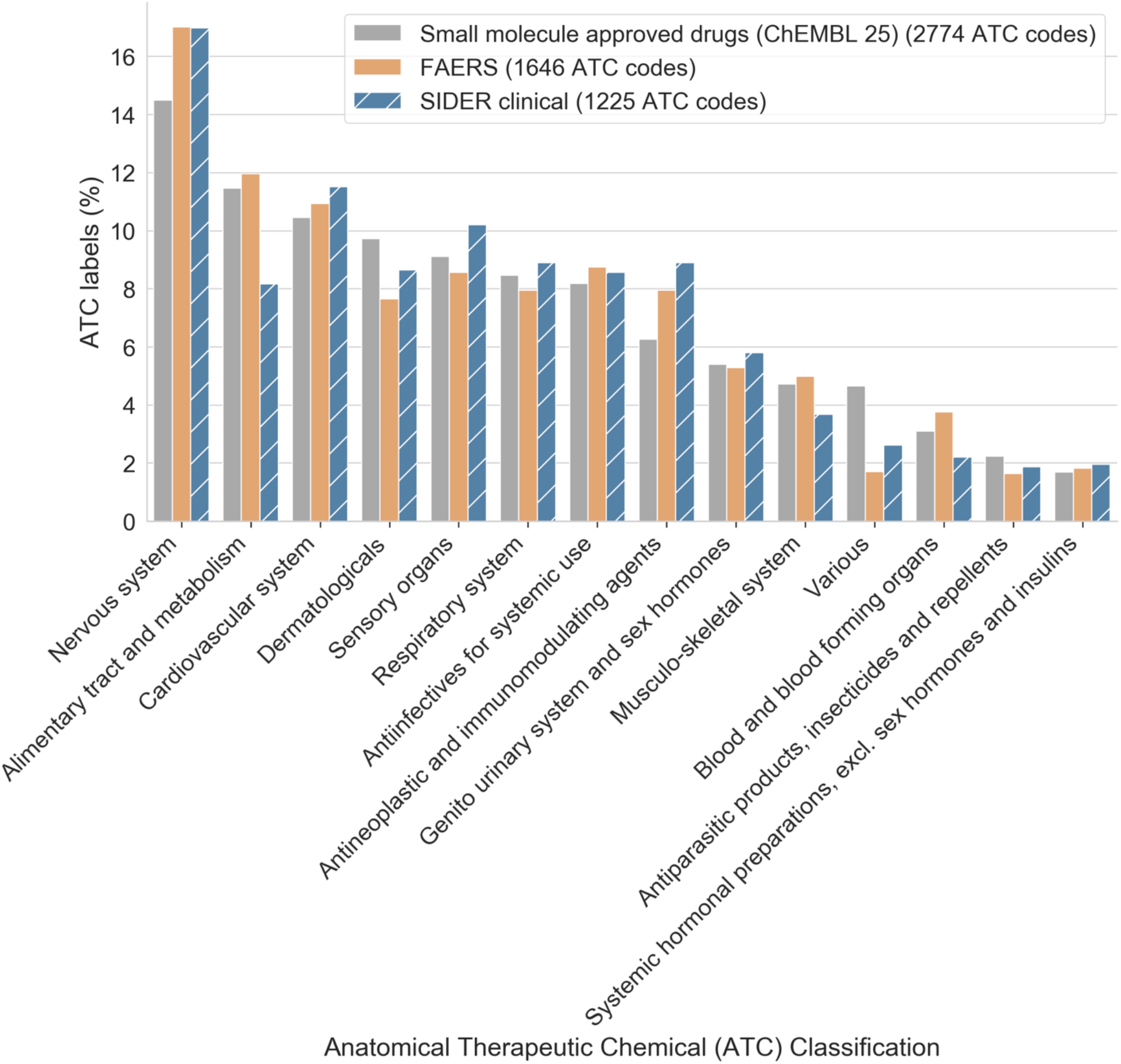
Drugs by their first level classification in the Anatomical Therapeutic Chemical (ATC) Classification, representing drug indications. If a drug has multiple ATC codes, all are included in the counts. The FAERS and SIDER datasets have a similar distribution of ATC labels as marketed small molecule drugs, showing they are largely representative of small molecule drug indications, although there are some differences.

**Fig. S2.**
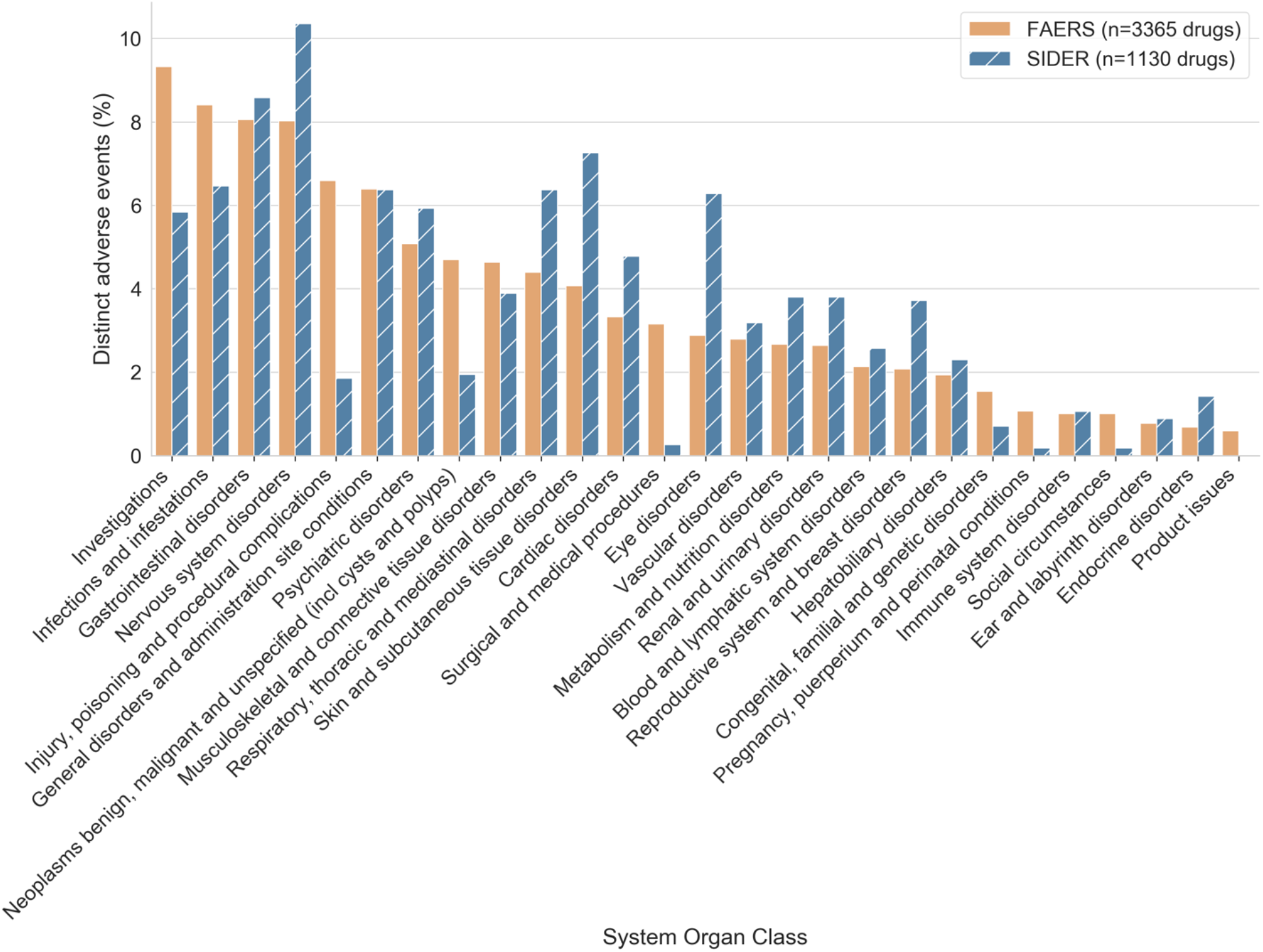
Diversity of AEs in the FAERS and SIDER datasets across System Organ Class. Each distinct AE is counted only once, even if associated with multiple drugs. There are differences in the types of AEs between FAERS and SIDER, for instance FAERS contains a greater diversity of events in the ‘Injury, poisoning and procedural complications’ class, whereas SIDER contains a relatively greater variety of Eye disorders.

**Fig. S3.**
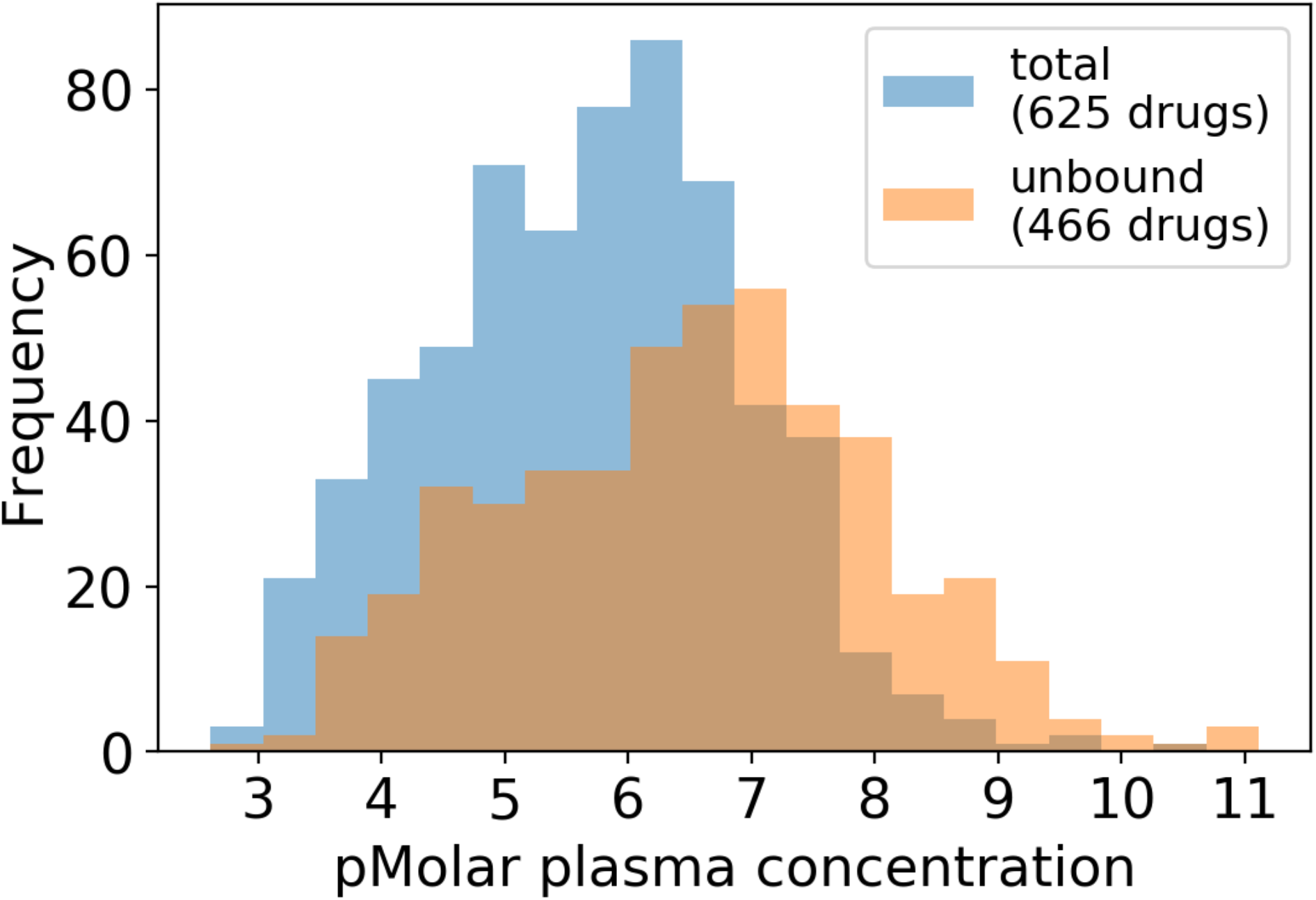
Distribution of total and unbound plasma concentrations compiled for the drugs in the FAERS and SIDER datasets. As expected, the unbound concentrations are lower, with a median of pMolar concentration of 6.6 and standard deviation (STD) of 1.5, compared to the total concentrations (median 5.8 and STD 1.3).

**Fig. S4.**
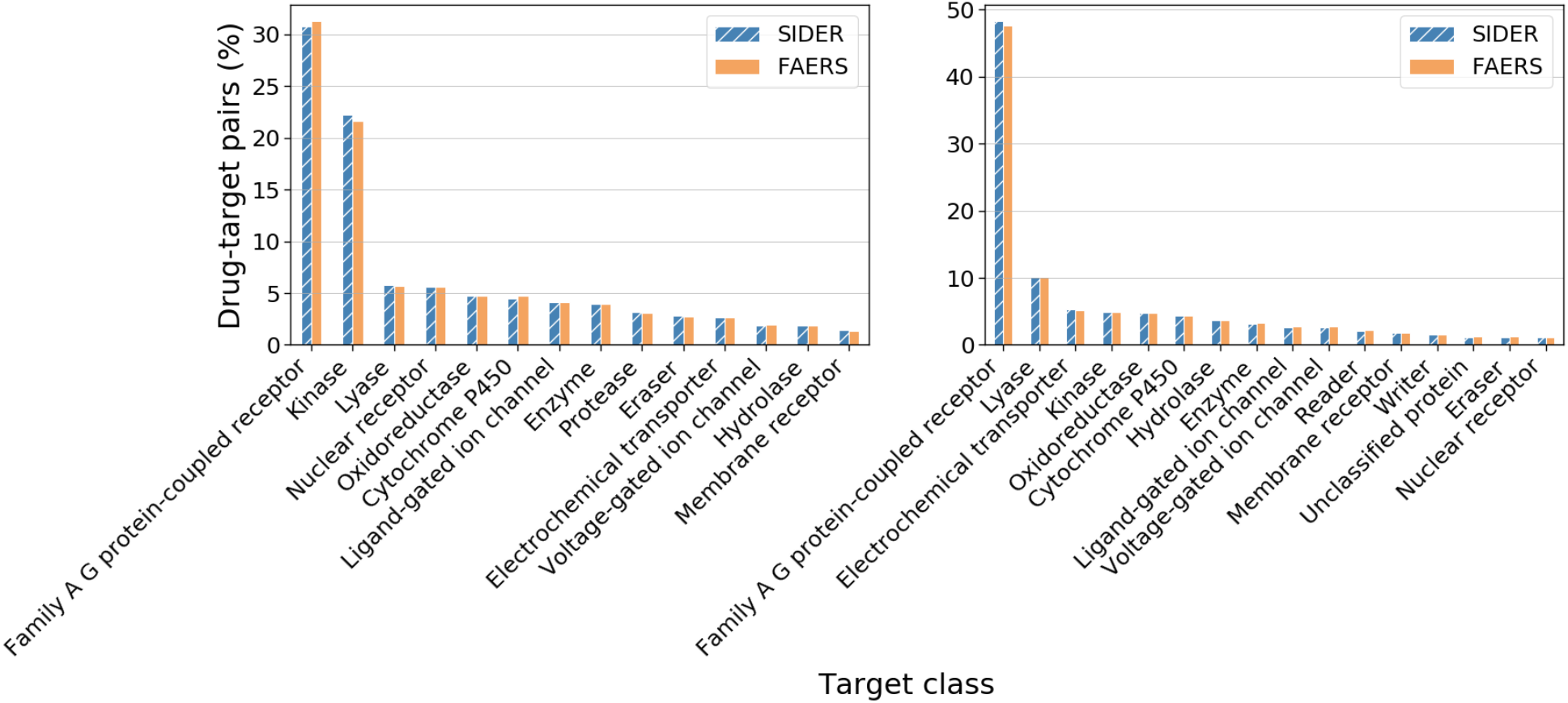
Distribution of target classes of the all drug-target bioactivity pairs (active and inactive) that are included when using a constant pChEMBL cut-off ≥ 6 (left panel) versus the subset of drug-target pairs that have unbound plasma concentrations available (right panel). Target classes are shown up to the second level of the ChEMBL target hierarchy if available (e.g. lyase), otherwise only the first level (e.g enzyme). When using the plasma concentrations the proportion of Family A G protein-coupled receptor datapoints is higher at the expense of kinase datapoints compared to using the constant cut-off.

**Fig. S5.**
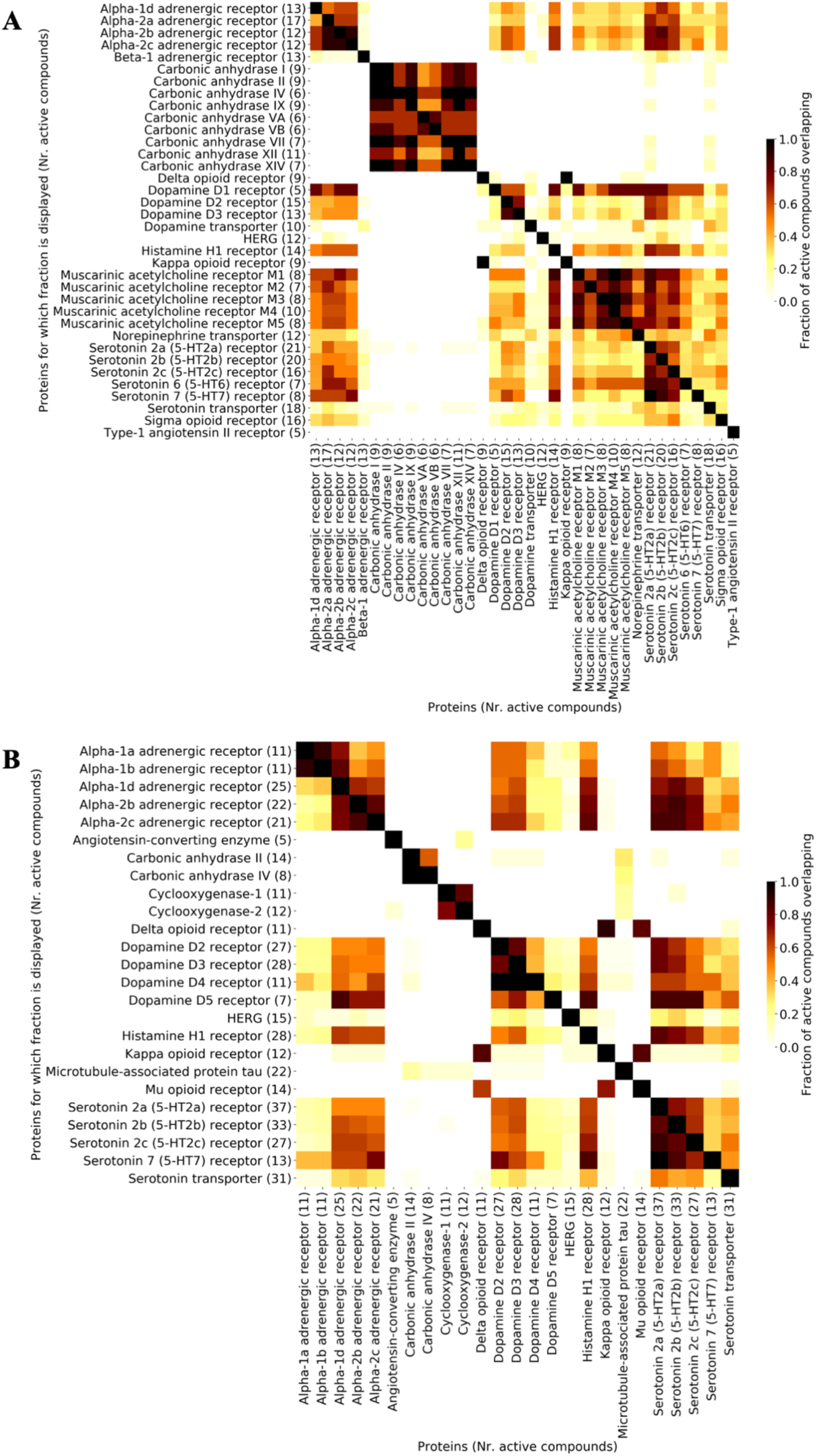
Fraction of active drugs, defined by the ratio of the *in vitro* bioactivity over the drug plasma concentration, shared between targets in the (**A**) SIDER and (**B**) FAERS dataset. The fraction corresponds to the target on the y-axis. The number of active compounds at each protein is shown in parentheses after the protein name. Targets often share active ligands, especially within target families, but also across families. In contrast, the carbonic anhydrases share ligands within the family but hardly so with other targets from other families.

**Table S1.**
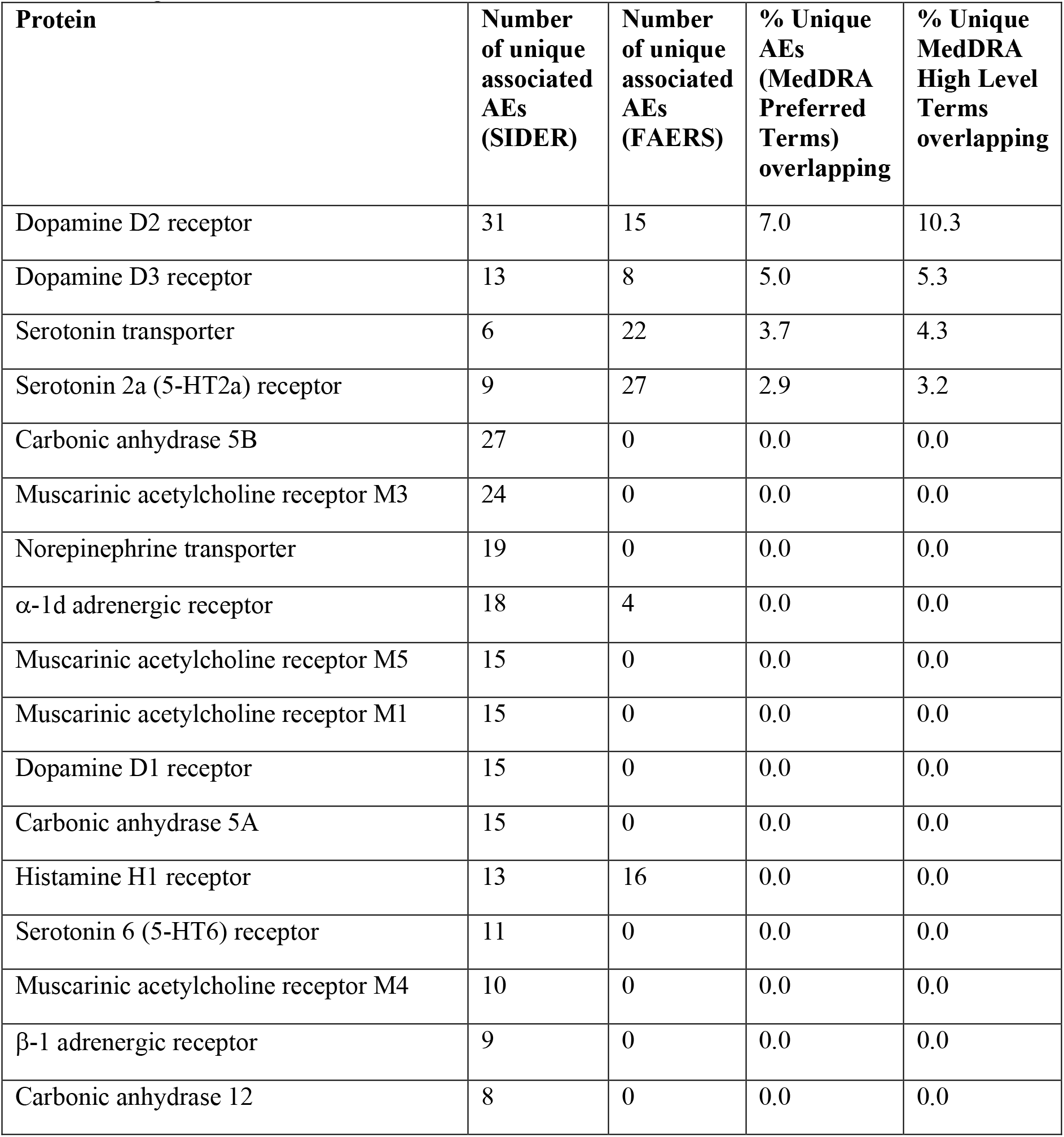
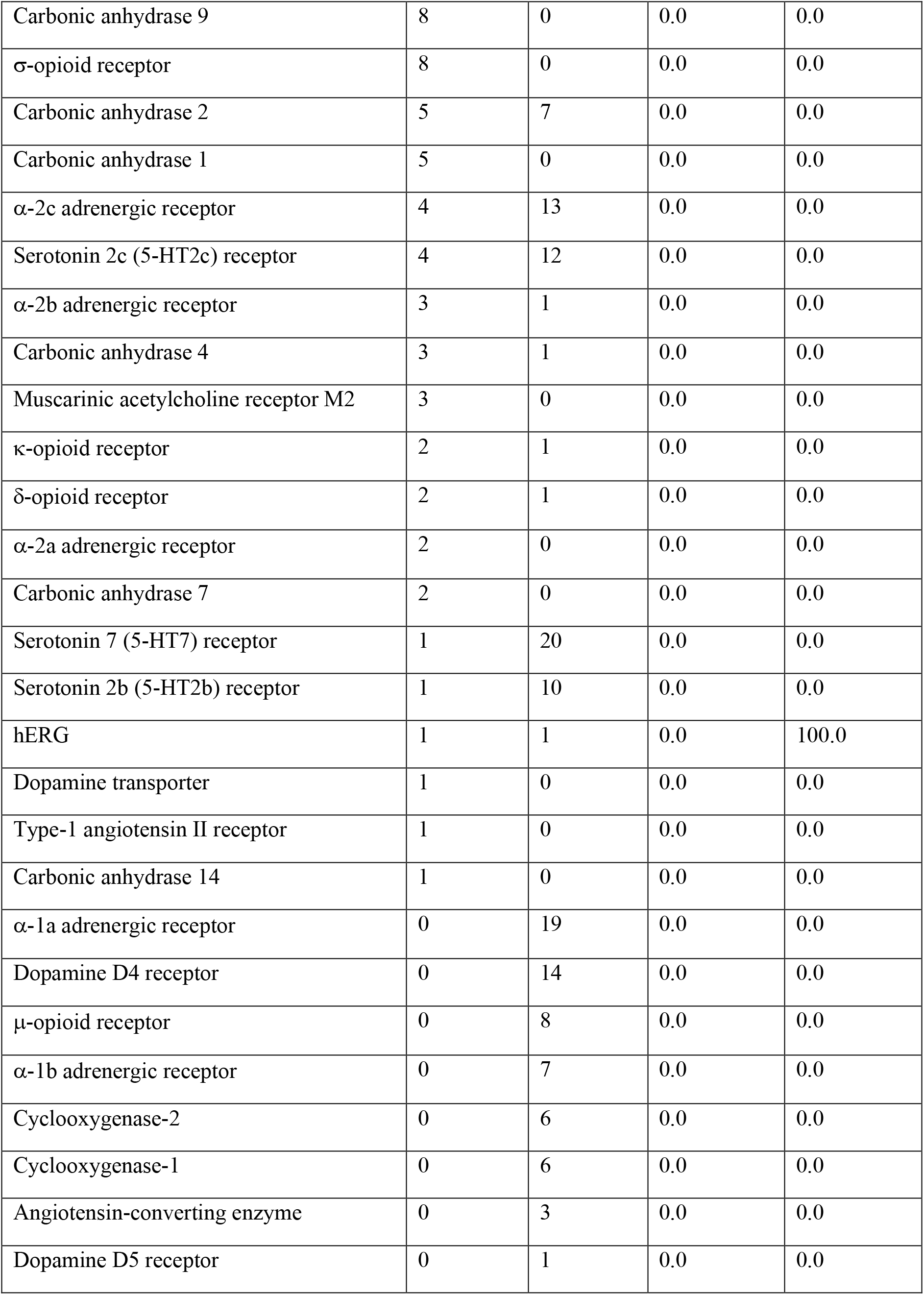
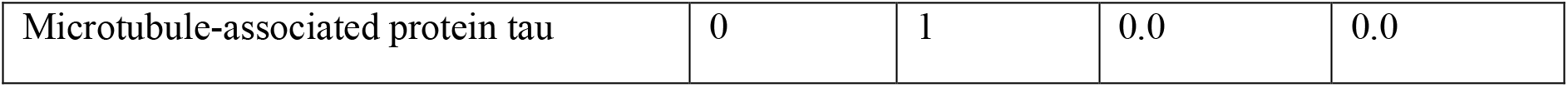
Targets with the number of significantly associated AEs in the SIDER and FAERS datasets and the percentage overlap between both sources. The majority of AEs associated with the same target is different in either dataset.

## Supplementary Materials

Data File S 1. Drug-AE relationships based on FAERS.

Data File S 2. Bioactivity data plus predictions used in the analysis.

Data File S 3. All positive target-AE combinations assessed for FAERS using the unbound plasma concentrations.

Data File S 4. All positive target-AE combinations assessed for SIDER using the unbound plasma concentrations.

Data File S 5. All positive target-AE combinations assessed for FAERS using the constant pChEMBL cut-off.

Data File S 6. All positive target-AE combinations assessed for SIDER using the constant pChEMBL cut-off.

Data File S 7. Share of measured versus predicted bioactivities per target for SIDER. Data File S 8. Share of measured versus predicted bioactivities per target for FAERS. Data File S 9. Extracted total drug plasma concentrations with references.

Data File S 10. Computed median unbound plasma concentrations used in the analysis.

Data File S 11. Previously reported safety target associations extracted and mapped to MedDRA terms (PT).

Data File S 12. Previously reported safety target associations extracted and mapped to MedDRA terms (HLT).

